# Massively parallel fitness profiling reveals multiple novel enzymes in *Pseudomonas putida* lysine metabolism

**DOI:** 10.1101/450254

**Authors:** Mitchell G. Thompson, Jacquelyn M. Blake-Hedges, Pablo Cruz-Morales, Jesus F. Barajas, Samuel C. Curran, Christopher B. Eiben, Nicholas C. Harris, Veronica T. Benites, Jennifer W. Gin, William A. Sharpless, Frederick F. Twigg, Will Skyrud, Rohith N. Krishna, Jose Henrique Pereira, Edward E. K. Baidoo, Christopher J. Petzold, Paul D. Adams, Adam P. Arkin, Adam M. Deutschbauer, Jay D. Keasling

## Abstract

Despite intensive study for 50 years, the biochemical and genetic links between lysine metabolism and central metabolism in *Pseudomonas putida* remain unresolved. To establish these biochemical links, we leveraged Random Barcode Transposon Sequencing (RB-TnSeq), a genome-wide assay measuring the fitness of thousands of genes in parallel, to identify multiple novel enzymes in both L- and D-lysine metabolism. We first describe three pathway enzymes that catabolize L-2-aminoadipate (L-2AA) to 2-ketoglutarate (2KG), connecting D-lysine to the TCA cycle. One of these enzymes, PP_5260, contains a DUF1338 domain, a family with no previously described biological function. Our work also identified the recently described CoA independent route of L-lysine degradation that metabolizes to succinate. We expanded on previous findings by demonstrating that glutarate hydroxylase CsiD is promiscuous in its 2-oxoacid selectivity. Proteomics of select pathway enzymes revealed that expression of catabolic genes is highly sensitive to particular pathway metabolites, implying intensive local and global regulation. This work demonstrates the utility of RB-TnSeq for discovering novel metabolic pathways in even well-studied bacteria, as well as a powerful tool for validating previous research.

**Importance:** *P. putida* lysine metabolism can produce multiple commodity chemicals, conferring great biotechnological value. Despite much research, connecting lysine catabolism to central metabolism in *P. putida* remained undefined. Herein we use Random Barcode Transposon Sequencing to fill in the gaps of lysine metabolism in *P. putida*. We describe a route of 2-oxoadipate (2OA) catabolism in bacteria, which utilizes DUF1338 containing protein PP_5260. Despite its prevalence in many domains of life, DUF1338 containing proteins had no known biochemical function. We demonstrate PP_5260 is a metalloenzyme which catalyzes an unusual 2OA to D-2HG decarboxylation. Our screen also identified a recently described novel glutarate metabolic pathway. We validate previous results, and expand the understanding of glutarate hydroxylase CsiD by showing can it use either 2OA or 2KG as a cosubstrate. Our work demonstrates biological novelty can be rapidly identified using unbiased experimental genetics, and that RB-TnSeq can be used to rapidly validate previous results.

## Introduction

*Pseudomonas putida* is an ubiquitous saprophytic soil bacterium and is a model organism for bioremediation (1). Interest in utilizing *P. putida* KT2440 as a chassis organism for metabolic engineering has recently surged due to the existence of well-established genetic tools and its robust metabolism of aromatic compounds that resemble lignin hydrolysis products (2–4). As lignin valorization remains essential for the economic feasibility of cellulosic bioproducts, a nuanced and predictable understanding of *P. putida* metabolism is highly desirable (5).

Although its aromatic metabolism has garnered much attention, the lysine metabolism of *P. putida* has also been rigorously studied for over fifty years (6). An understanding of lysine metabolism has had biotechnological value, as it has been used to produce glutarate, 5-aminovalerate (5AVA), as well as valerolactam in *P. putida* and in the other bacteria (7–10). Our current understanding of lysine catabolism however, remains incomplete. In particular, the connection between D-lysine metabolism and central metabolism in *P. putida* is unclear and has not been fully characterized.

*P. putida* employs bifucating pathways to catabolize lysine, separately metabolizing the L- and D-isomers (Figure S1a) (11). The L-lysine degradation pathway proceeds to glutarate, which can then be either be degraded to acetyl-CoA via a glutaryl-CoA intermediate, or to succinate without a CoA bound intermediate (Figure S1a) (9). The final steps of D-lysine catabolism remain more elusive. The initial steps of D-lysine catabolism are well described, but the genetic basis stops at 2AA (12). Furthermore, ^13^C labeling experiments by Revelles et al. demonstrated a putative metabolic connection between the D- and L-lysine pathways at 2AA (11). Subsequent steps to central carbon metabolism have never been fully validated. (6, 11–13). Given the importance of lysine metabolism, and recent availability of high-throughput genetic tools, we sought to identify the missing steps in D-lysine metabolism that have remained despite 50 years of research.

Random barcode transposon sequencing (RB-TnSeq) is a genome-wide approach that measures the importance of each gene to growth (or fitness) in a massively parallel assay (14). RB-TnSeq can identify phenotypes for thousands of previously uncharacterized genes (14, 15), including the levulinic acid degradation pathway in *P. putida* KT2440 (16). In this study, we applied RB-TnSeq to uncover multiple novel genes implicated in L- and D-lysine metabolism in *P. putida*. We first describe a three enzyme route connecting L-2AA to 2KG (Figure S1B). Within this pathway, D-lysine metabolism connects to central metabolism through a 2HG intermediate, which is directly produced from 2OA in a reaction catalyzed by a DUF1338-containing protein. This protein family, widely distributed across many domains of life, previously had no known function. Subsequently, we further characterize the glutarate hydroxylase CsiD, by demonstrating its 2-oxoacid promiscuity during the hydroxylation of glutarate. Finally, we show the expression of all newly discovered enzymes changes significantly in response to specific metabolites within the two catabolic pathways.

## Results

### Identification of lysine catabolism genes via RB-TnSeq

To identify mutants defective in lysine catabolism in *P. putida* KT2440, an RB-TnSeq library of this bacterium (16) was grown on minimal medium supplemented with either D-lysine or L-lysine as the sole carbon source. To evaluate whether D-lysine metabolism was required for the metabolism of other downstream metabolites of L-lysine, the library was also grown on 5AVA. As a control, we also grew the library on glucose. Fitness was calculated as the log_2_ ratio of strain and gene abundance at the end of selective growth relative to initial abundance (14). Fitness profiling revealed 39 genes with significant fitness values of less than −2 for 5AVA, D-lysine, or L-lysine, and no less than −0.5 fitness for glucose (Figure 1a, Supplementary Table 1). Within this set, 10 of the 12 known lysine degradation genes were identified, with the exception of the two enzymes in the CoA-dependent route of glutarate degradation (*gcdH* and *gcdG*), which both had significant fitness values (*t* < −4) but whose fitness was greater than −2. Instead, we identified the recently-characterized genes involved in the CoA independent pathway (*csiD* and *lghO*) (9).

**Figure 1:**
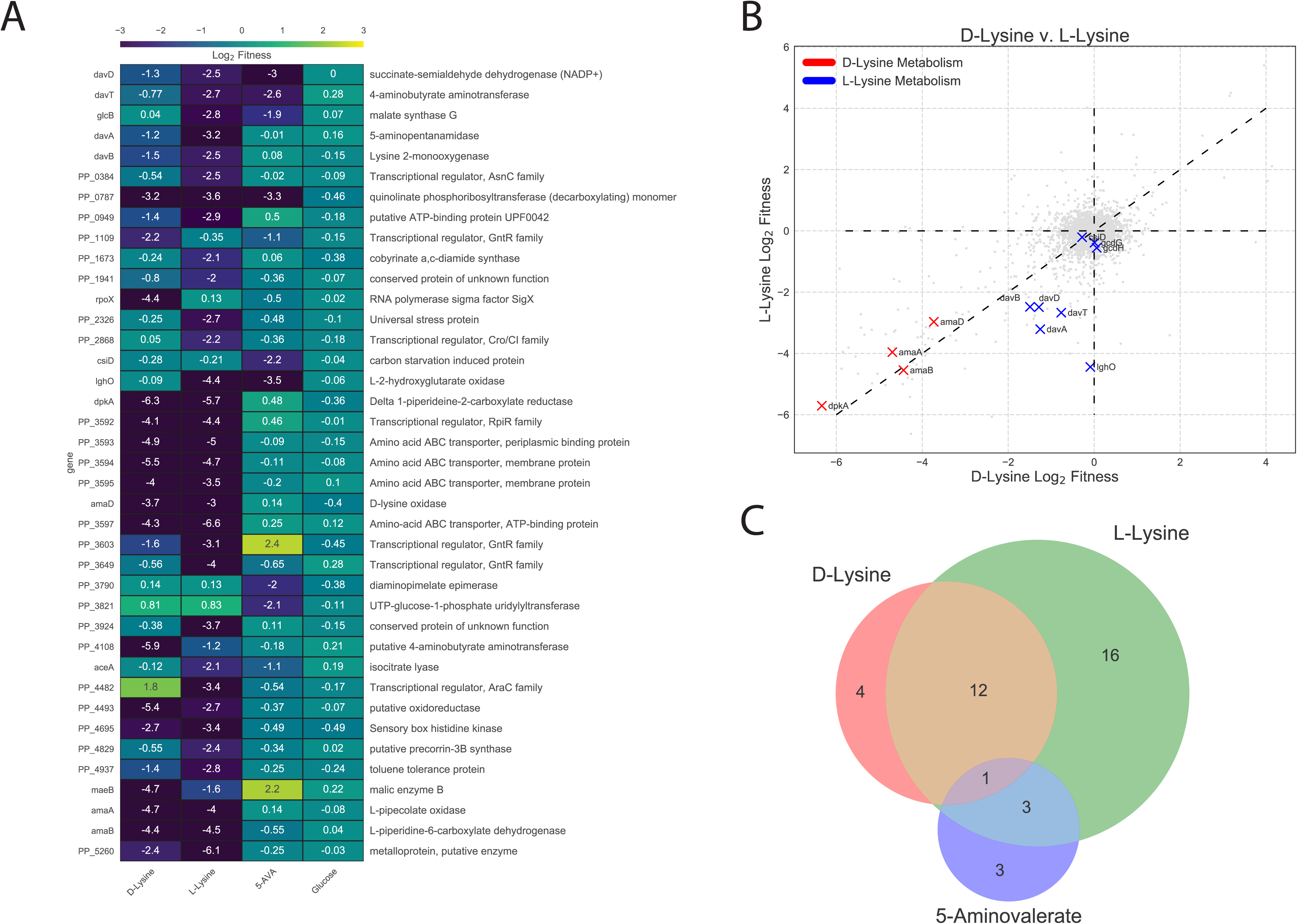
Results of RB-TnSeq screen. A) Genes that showed less than −2 log_2_ fitness on either D-lysine, L-lysine, or 5AVA but showed no less than −0.5 log_2_ fitness defect when grown on glucose. B) Plot of genome wide fitness values of libraries grown on either L-lysine or D-lysine. Genes encoding for enzymes known to be involved in D-lysine metabolism are shown in red, while those known to be involved in L-lysine metabolism are shown in blue. C) Venn diagram of genes with significant fitness defects when grown on either D-lysine, L-lysine, or 5AVA.

The fitness data corroborated previous work showing a functional D-lysine pathway is required for L-lysine catabolism (6, 11). None of the known L-lysine catabolic genes showed fitness defects for growth on D-lysine, but transposon insertions in all previously-identified D-lysine genes showed negative fitness scores when grown on L-lysine (Figure 1b). No known D-lysine catabolic enzymes showed fitness defects when grown on 5AVA, suggesting the D-lysine dependence of L-lysine catabolism may only occur for early catabolic steps (Figure 1c).

In addition to catabolic enzymes, lysine transporters and multiple transcriptional regulators were identified (Figure 1a). The putative lysine amino acid ABC transporter system (PP_3593, PP_3394, and PP_3395) showed significant fitness defects when grown with either isomer of lysine. Some of the transcriptional regulators were located near known catabolic or transport enzymes (PP_0384, PP_3592, and PP_3603), while others were not clustered with any obviously related genes (PP_1109, PP_2868, PP_3649, and PP_4482). Two known global regulators were identified in our screen: *cbrA* (PP_4695), a histidine kinase sensor that showed fitness defects on both lysine isomers, and the alternative sigma factor *rpoX* (PP_2088) which only had fitness defects when grown on D-lysine.

Additionally there were 15 genes which, when disrupted, displayed fitness advantages greater than 2 on 5AVA, D-lysine, or L-lysine and less than 0.5 fitness when grown on glucose. This positive fitness value indicates these mutations confer a competitive advantage compared to other strains when grown on these carbon sources. Most striking amongst these genes were the sigma factor *rpoS* and the LPS export system (PP_1778/9), which when disrupted, both displayed fitness benefits on all three non-glucose carbon sources (Figure S2).

Only one gene (PP_0787, a quinolinate phosphoribosyltransferase) showed fitness defects on all three non-glucose carbon sources (Figure 1c). However, disruption of PP_0787 also showed a significant fitness defect when grown on levulinic acid, suggesting it is unlikely to be uniquely important to lysine metabolism (16). Only 3 genes shared fitness defects between 5AVA and L-lysine (*davT*, *davD*, and *lghO*), all of which have been previously implicated in 5AVA metabolism (Figure 1c) (9).

### PP_4108 is a L-2AA aminotransferase

In humans and other animals, L-lysine degradation proceeds through a 2AA intermediate, which a transaminase converts to 2OA (9, 11, 17). Yet, no such transaminase has been identified in *P. putida*. We identified a candidate aminotransferase, PP_4108, for which gene inactivation showed a significant growth defect on D-lysine (−5.9) and a relatively minor defect on L-lysine (−1.2). To corroborate our RB-TnSeq fitness data, we constructed a deletion mutant of PP_4108 that failed to grow in a plate reader assay on 10 mM DL-2AA (Figure 2a). The mutant showed a severe growth defect on 10 mM D-lysine and an increased lag time when grown on 10 mM L-lysine (Figure S3).

**Figure 2:**
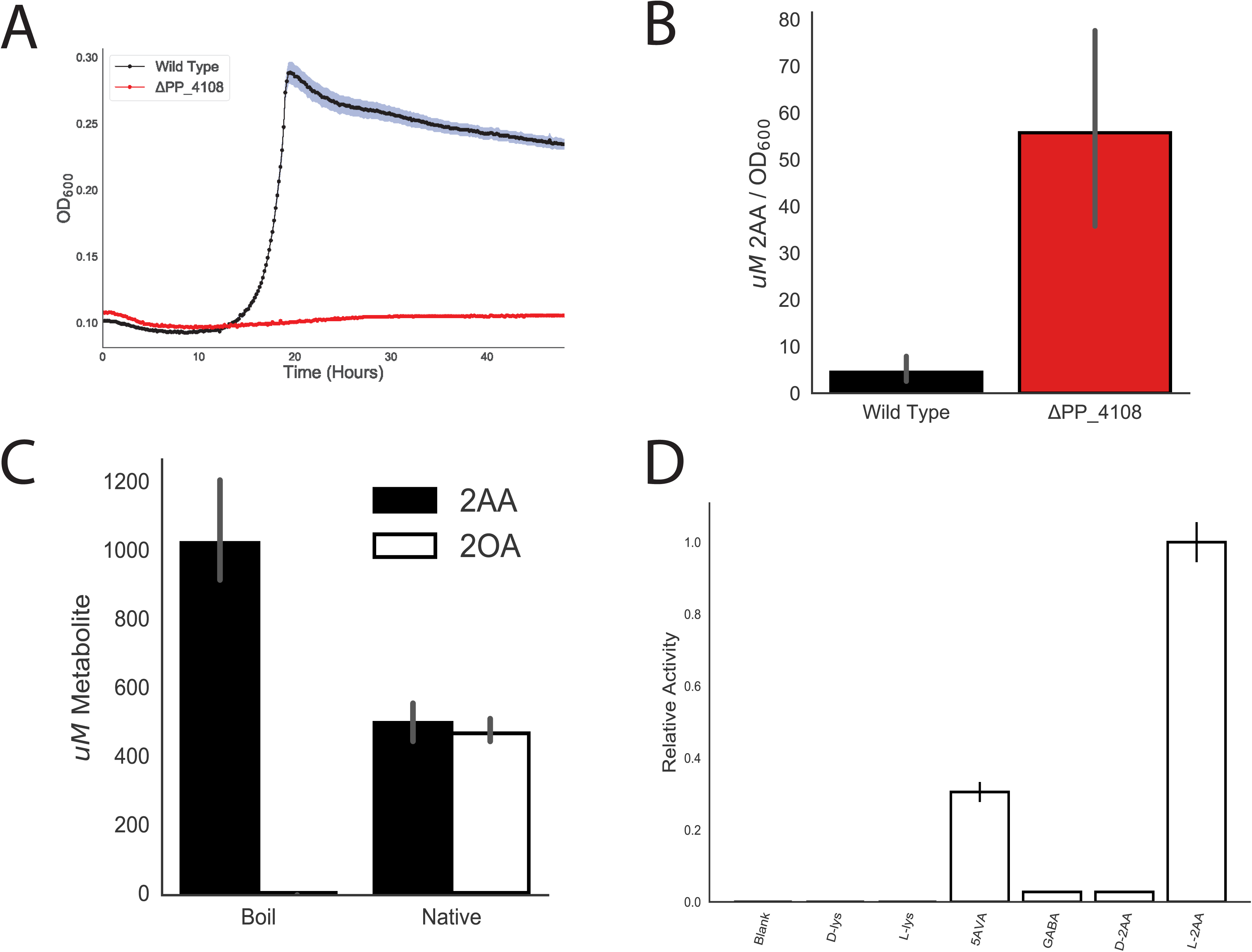
Identification of PP_4108 as a L-2AA aminotransferase. A) Growth of wild-type KT2440 and PP_4108 mutant on 2AA as a sole carbon source. Shaded area represents 95% confidence interval (CI), n=3. B) *In vivo* accumulation of 2AA in wild-type KT2440 and a PP_4108 mutant after 12 hours of growth on minimal medium supplemented with 10 mM glucose and 10 mM D-lysine. Bars represent 1og_10_ transformed spectral counts, error bars show 95% CI, n=3. C) *In vitro* transamination reactions of PP_4108 with 2KG as an amino acceptor. Bars represent µM metabolite concentration of either 2OA (black) or 2AA (white) in either boiled or native protein reactions. Error bars show 95% CI, n=3. D) *In vitro* transaminations of PP_4108 incubated with different possible amino donors and 2KG as acceptor. Bars represent relative activity of enzyme standardized to L-2AA after 16 hour incubation. Error bars show standard deviation, n=2.

To further validate this hypothesis, the ΔPP_4108 strain was subjected to metabolomics analysis to monitor the accumulation of its expected substrate, 2AA, when grown on glucose and D-lysine. After 12 hours of growth on minimal media supplemented with 10 mM each of glucose and D-lysine, the PP_4108 deletion strain showed a 6.3-fold increase (*p* = 0.00016) in normalized intracellular 2AA concentration compared to WT (Figure 2b). Next PP_4108 was expressed and purified from *E. coli* for biochemical characterization. After purified enzyme incubation with DL-2AA, 2KG, and pyridoxal phosphate (PLP) for 16 hours, the reaction mixture was analyzed with LC-TOF. The expected product, 2OA, was detected in the enzymatic reaction but not in a boiled enzyme control, confirming PP_4108 as a transaminase that converts 2AA to 2OA (Figure 2c). As many transaminases have broad substrate specificity (18), we also probed the substrate range of PP_4108 using a colorimetric assay for glutamate, a stoichiometric product of the transamination reaction (Figure 2d). The enzyme was most active on L-2AA, and only showed 2.8% relative activity (*p* = 0.0057) on its enantiomer, D-2AA. This specificity for the L-2AA isomer may explain why only 50% of the DL-2AA was transformed in the previous experiment (Figure 2c). No activity was observed on either lysine isomer; however, the enzyme had slight activity towards 4-aminobutyrate/γ-aminobutyrate (GABA) (2.8% relative activity, *p* = 0.0057) and moderate activity on 5AVA (30.5% relative activity, *p* = 0.0139). Over shorter time scales PP_4108 had no activity on any substrate expect L-2AA (Figure S3c). These results suggest *P. putida* KT2440 metabolizes D-lysine to L-2AA, which is then converted to 2OA by the transaminase PP_4108.

### PP_5260 is a novel DUF1338 family enzyme that catalyzes the conversion of 2OA to 2HG

Early work proposed 2OA is converted to 2KG via a 2HG intermediate (13, 19), while later results suggested a direct conversion of 2OA to glutarate (11). Either route likely requires a decarboxylation of 2OA, so we initially searched for decarboxylases within our dataset. Our fitness data on either lysine isomer revealed no obvious decarboxylases or enzymes likely to contain a thiamine pyrophosphate (TPP) cofactor, which are commonly employed by decarboxylases. However, a gene near other D-lysine catabolic genes in the *P. putida* genome, PP_5260, showed a significant fitness defect. A ΔPP_5260 strain was unable to grow on either isomer of lysine verifying its importance in lysine degradation (Figure 3a).

**Figure 3:**
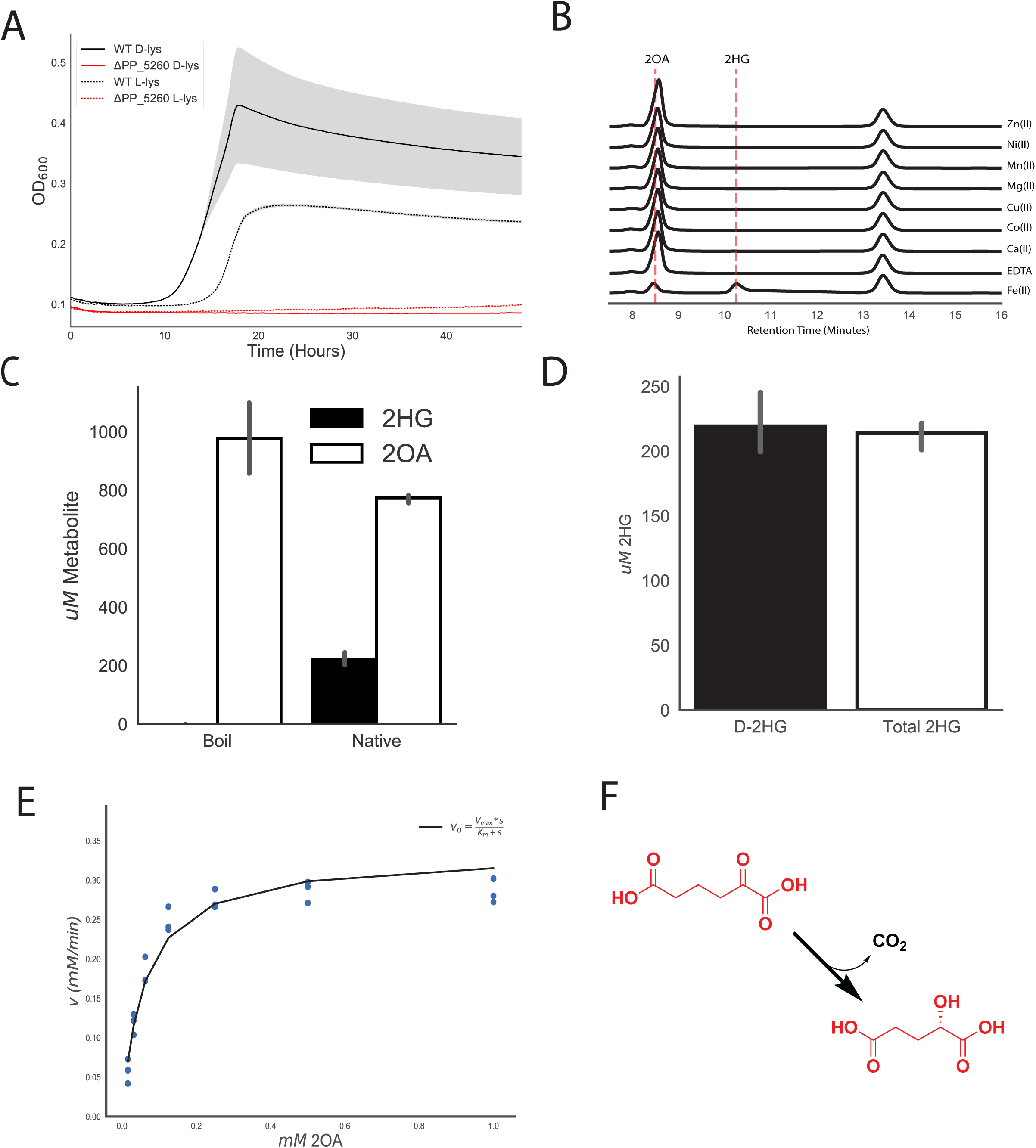
Identification of ydcJ (PP_5260) as a 2OA decarboxylase/hydroxylase. A) Growth of wild-type (black) and PP_5260 mutant (red) on D-lysine (line) or L-lysine (dashed line) as a sole carbon source. Shaded area represents 95% CI, n=3. B) HPLC traces of *in vitro* reactions run with apo PP_5260 with exogenous metals added at 50 µM. Retention times for 2OA and 2HG are shown by vertical dashed lines. Metal or EDTA control is indicated to the right of traces. C) *In vitro* assay of 2OA conversion to 2HG by purified PP_5260 protein analysed via LC-TOF. 2OG in white, 2HG in black. D) *In vitro* assay of purified PP_5260 protein with 2OA as substrate. Black bar represents concentration of D-2HG measured by enzyme coupled assay. White bar represents total 2HG concentration as measured by LC-TOF. Error bars represent 95% CI, n=3. E) Initial velocity of reaction catalyzed by PP_5260 as a function of 2OA concentration. Blue dots represent individual measurements, while the black fit line shows a Michaelis-Menten fit. F) Chemical reaction catalyzed by PP_5260, 2OA is decarboxylated to D-2HG.

PP_5260 belongs to the DUF1338 protein family (http://pfam.xfam.org/family/PF07063). Although several unpublished crystal structures of DUF1338 domain containing proteins have been deposited into the Protein Data Bank, their biological function remains elusive. However, these structures combined with protein sequence alignments suggest a putative metal binding site is conserved throughout the DUF1338 family. As we hypothesized PP_5260 serves as the missing decarboxylase in D-lysine metabolism, we purified the enzyme for biochemical analysis. Enzymatic activity on 2OA was probed and analyzed via LC-TOF. After incubation of 2OA with PP_5260, we observed a ∼92% (*p*=0.00034) reduction in the abundance of 2OA, whereas no 2OA was consumed in a boiled enzyme control or enzyme treated with EDTA confirming it to be a metalloenzyme (Figure S4a). Initially we believed the product would be either glutarate or glutarate semialdehyde, however neither of these was detected in the reaction. Early biochemical work suggested 2HG as a potential intermediate in pipecolate metabolism (19), and when the enzymatic product was compared to a racemic 2HG standard they shared the same mass, retention time and mass-to-charge ratio (Figure S4b), as well identical isotopic distributions of [M-H] peaks in the mass spectra (Figure S4c).

To identify the metal cofactor, the enzyme was dialyzed against EDTA to remove metals, and individual divalent metals were added back. Only the addition of Fe(II) restored enzymatic activity as measured by HPLC (Figure 3b). Subsequent reactions quenched after 5 minutes showed 200 uM of 2HG formed and 800 uM 2OA remaining, demonstrating 1:1 2OA to 2HG reaction stoichiometry (Figure 3c). Whether the product of the PP_5260 reaction is either L-2HG or D-2HG is critical to understanding the eventual fate of D-lysine, as *lghO* is specific for L-2HG (Figure S1a). An enzyme coupled assay specific for the detection of D-2HG was used to assess the stereochemistry of the PP_5260 product. Standard curves of D-2HG and L-2HG showed that the assay was only responsive to D-2HG (Figure S4d). The concentration of *in vitro* reactions of PP_5260 were then measured by both LC-TOF as well as the enzyme coupled assay, revealing all 2HG as the D-isomer (Figure 3d).

Kinetic parameters of PP_5260 were determined using an enzyme-coupled assay to spectrophotometrically measure CO_2_ evolution via NADH oxidation (20). PP_5260 displayed a V_max_ of 0.33 mM/min (+/− 0.08 mM), a K_m_ of 0.06 mM (+/− 0.03 mM), and a K_cat_ of 330 m^−1^ using 2OA as a substrate. Taken together these results reveal that PP_5260 is novel Fe(II) dependent decarboxylase that converts 2OA to D-2HG (Figure 3f), a chemical reaction not previously observed in nature.

### DUF1338 proteins are a widely distributed enzyme family with a putative conserved role in amino acid catabolism

After functional characterization of PP_5260, we use phylogenomics to propagate the annotation and further explore the biological role of DUF1338 proteins found in other organisms. We found that DUF1338 proteins are widely distributed across the tree of life, with homologs of PP_5260 found in plants and green algae (22), fungi, and bacteria, though they were not found in animals or archaea (Figure 4a). Homologs are widely distributed amongst bacteria, with the Firmicutes being a notable exception. PP_5260 homologs within the plant group Streptophyta, as well as bacterial groups Actinobacteria, Cyanobacteria, and Bacteroidetes formed monophyletic clades, while homologs from other taxonomic groups were not monophyletic (Figure 4a). DUF1338 homologs are found in bacteria important to biotechnology (*Corynebacterium glutamicum*), the environment (*Nostoc puncitforme*), and medicine (*Yersinia pestis*, *Mycobacterium tuberculosis*, *Burkholderia pseudomallei*).

**Figure 4:**
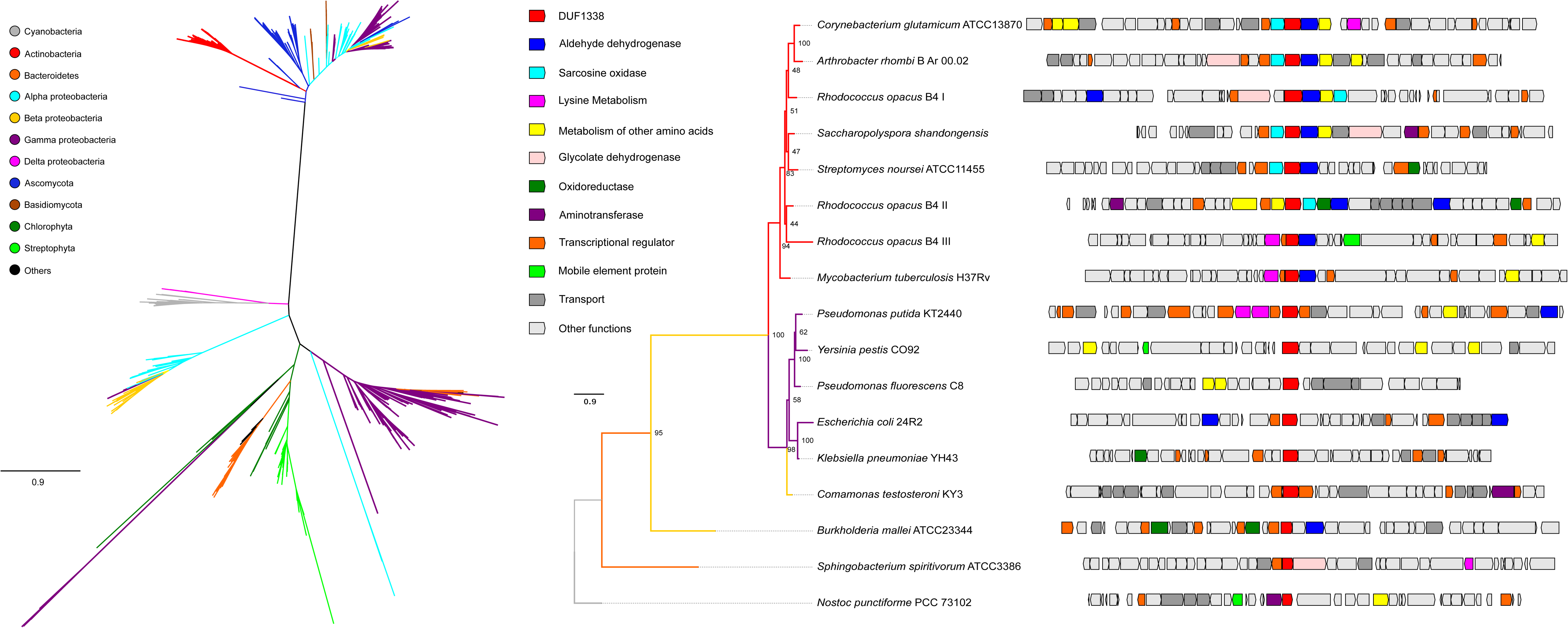
Phylogenomics of the DUF1338 enzyme family. A) Phylogenetic relationships among DUF1338 homologs and their distribution among major phyla. Branches in the tree are colored by phylum. DUF1338 is found in most bacterial phyla as well as in plants and fungi. Non-monophyletic clades suggest pervasive horizontal gene transfer events in the family. B) Phylogenomics of selected DUF1338 homologs in bacteria. The phylogeny in the left shows the phylogenetic relationships between selected homologs, the branches have been colored according to their adscription to a given phylum and the support values are shown at the nodes. The boxes in the right represent the gene neighborhood for each homolog. The genes have been colored to represent their annotated functions.

Publicly available fitness data show both *Pseudomonas fluorescens* FW300-N2C3 and *Sinorhizobium meliloti* PP_5260 homologs have L-lysine specific defects when interrupted (15). Genomic contexts within other bacteria suggest many DUF1338-containing enzymes may be involved in lysine or other amino acid metabolism (Figure 4b). Within the Actinobacteria DUF1338 homologs are often found adjacent to sarcosine oxidases, aldehyde dehydrogenases, and transaminases implying an additional catabolic amino acid function. In both the oleaginous bacterium *Rhodococcus opacus* B4 and *M. tuberculosis*, DUF1338 homologs are found next to predicted L-lysine aminotransferases, suggesting an ancestral homolog functioned in lysine catabolism. Interestingly, the *R. opacus* B4 genome has three DUF1338 homologs, only one of which contains genes predicted to be specific to lysine catabolism. The other two gene neighborhoods are similar in their functional content, mainly differing by containing an oxidoreductase or glycolate dehydrogenase, either of which may perform the same biochemical function. In Alphaproteobacteria, Betaproteobacteria, and Cyanobacteria, the presence of aldehyde dehydrogenases, oxidoreductases, glycolate dehydrogenases, and aminotransferases implies a metabolic function similar to PP_5260.

### PP_4493 putatively oxidizes D-2HG to 2KG and connects D-lysine to central metabolism

In the CoA independent route of glutarate metabolism, LghO oxidizes L-2HG to 2KG, however this enzyme is highly specific towards the L-2HG isomer and showed no fitness defect on D-lysine in our RB-TnSeq data (Figure S1a). A putative FAD-dependent and 4Fe-4S cluster-containing glycolate dehydrogenase, PP_4493, did show fitness defects on both D-lysine and L-lysine (fitness scores of −5.4 and −2.7 respectively) (Figure 1a). Glycolate dehydrogenases are members of a larger family of enzymes that oxidize the alcohol group of an alpha-hydroxyacid to their corresponding alpha-ketoacid (Figure 5a). Therefore, we hypothesized PP_4493 could potentially oxidize a similar 2-hydroxyacid, 2HG, to the corresponding alpha-ketoacid, 2KG. Moreover, many PP_5260 homologs were located next to or near putatively annotated glycolate dehydrogenases in other bacteria, underscoring their potential metabolic link (Figure 4b). To confirm these hypotheses, we again constructed a deletion strain, *P. putida* ΔPP_4493, which could not grow on D-lysine as a sole carbon source (Figure 5b), and showed attenuated growth on L-lysine (Figure S5). Furthermore, when grown on 10 mM glucose and 10 mM D-lysine the mutant accumulated ∼500 uM 2HG normalized to optical density, whereas wild type *P. putida* did not accumulate any detectable 2HG (Figure 5c). Subsequent analysis of accumulated 2HG via a D-2HG specific detection kit revealed that this accumulated 2HG was indeed D-2HG (Figure 5c). These data and the conserved function and genomic context of glycolate dehydrogenases strongly suggest PP_4493 catalyzes the last step of L-2AA metabolism, oxidizing D-2HG to 2KG (Figure S1b).

**Figure 5:**
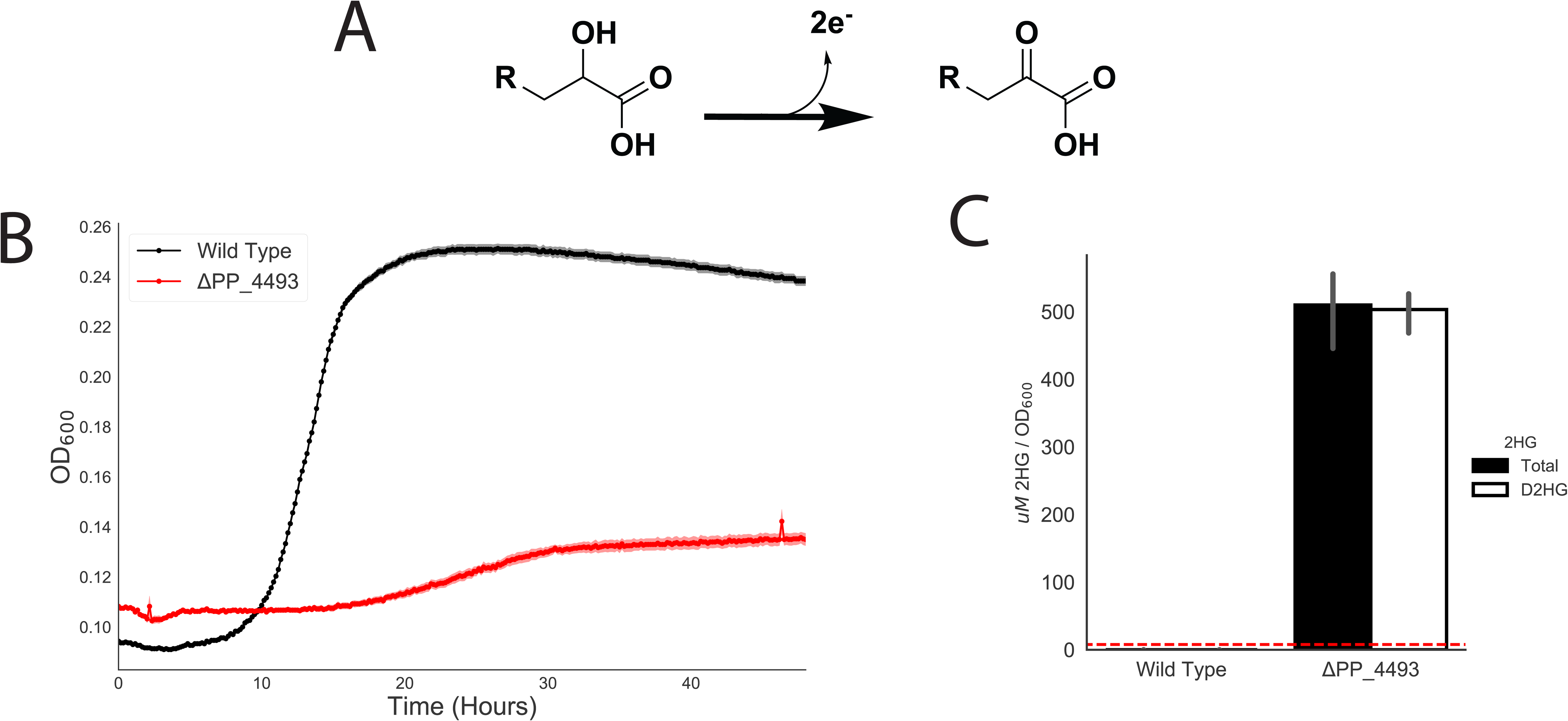
Identification of PP_4493 as a putative D-2HG dehydrogenase. A) General chemical reaction of a dehydrogenase converting a 2-hydroxyacid to a 2-ketoacid B) Growth of *P. putida* KT2440 and PP_4108 mutant on D-lysine as a sole carbon source. Shaded area represents 95% CI, n=3. C) *In vivo* accumulation of 2HG in wild-type KT2440 and a PP_4108 mutant after 12 hours of growth on minimal medium supplemented with 10 mM glucose and 10 mM D-lysine. White bar represents concentration of D-2HG measured by enzyme coupled assay. Black bar represent total 2HG concentration as measured by LC-TOF. Red line represents limit of detection of enzyme coupled assay for D-2HG. Bars represent 1og_10_ transformed spectral counts, error bars show 95% CI, n=3.

### CsiD is highly specific for glutarate hydroxylation but promiscuous in 2-oxoacid selectivity

During the initial preparation of this manuscript, Zhang et al. discovered a novel pathway of glutarate metabolism in *P. putida (9)*. They describe a cyclic reaction cascade wherein a novel 2KG-dependent non-heme Fe(II) oxygenase, PP_2909 (CsiD), hydroxylates glutarate to form 2HG and succinate using 2KG as a cosubstrate. PP_2910 (LghO), a putative L-2HG oxidase, then subsequently converts L-2HG to 2KG, regenerating the 2KG consumed in the initial reaction. These reactions result in the net incorporation of succinate into central metabolism (Figure S1). Our fitness results of the library grown on 5AVA also identified both *csiD* and *lghO*, in addition to the two enzymes from the CoA-dependent glutarate pathway, glutaryl-CoA ligase (*gcdG*) and glutaryl-CoA dehydrogenase (*gcdH*), mutants of which showed mild fitness defects when grown on 5AVA (Figure 6a). We also purified *csiD* and confirmed it hydroxylates glutarate in a 2KG-dependent manner (Figure S6a). HPLC analysis demonstrated that as glutarate was consumed, equimolar quantities of succinate and L-2HG were produced (Figure S6b). Additionally, a *csiD* deletion mutant showed increased lag time when grown on either L-lysine or 5AVA. By deleting the glutaryl-CoA ligase *gcdG*, and disrupting the CoA-dependent glutarate pathway, we completely prevented growth on 5AVA or L-lysine (Figure S6c). These results are in agreement with those found with Zhang et al (9).

**Figure 6:**
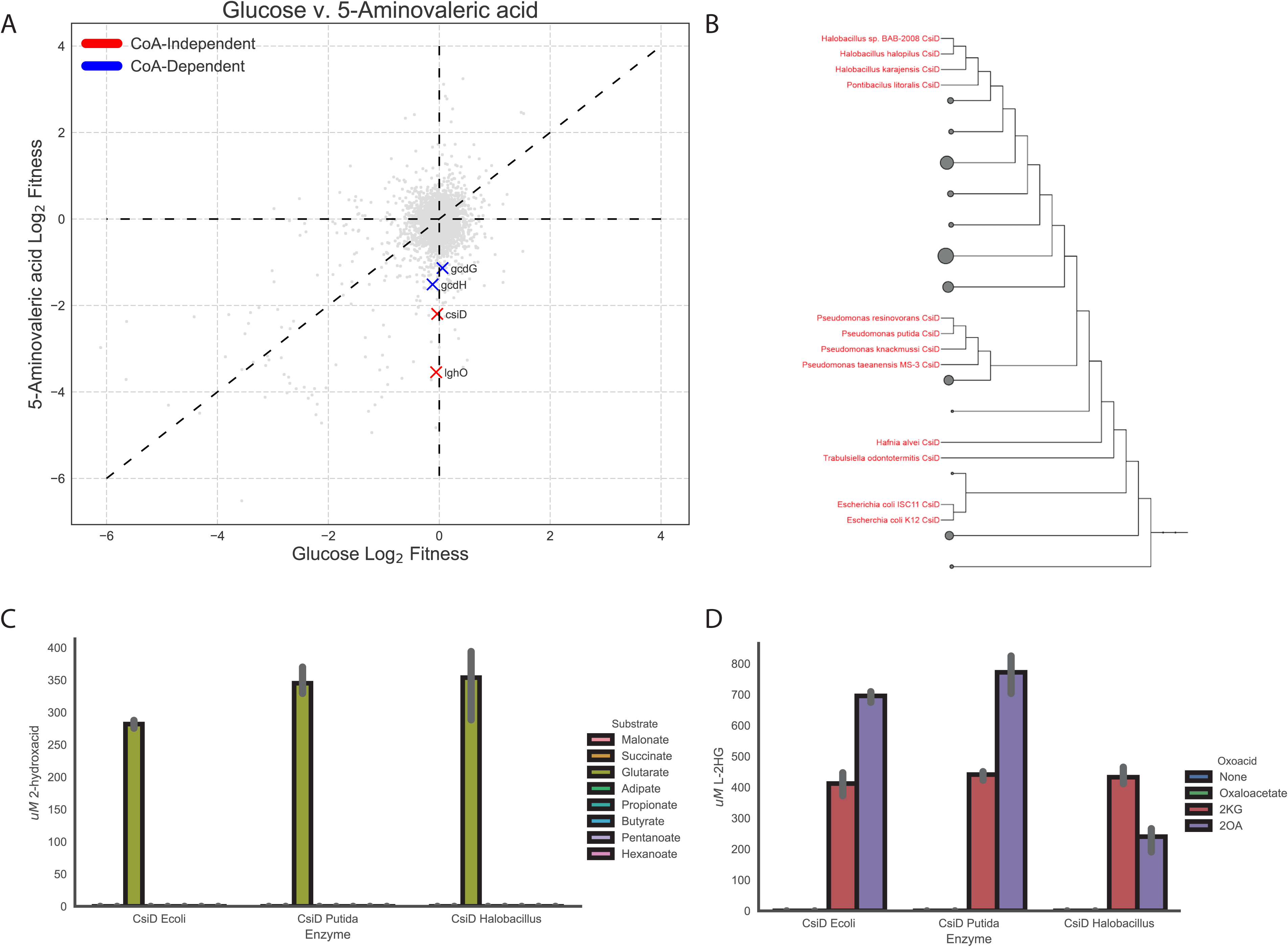
Role of CsiD in *P. putida* lysine metabolism. A) Plot of genome wide fitness values of libraries grown on either 5AVA or glucose. CoA-dependent glutarate degradation genes are shown in red, while those known to be involved succinate producing metabolism are shown in blue. B) Phylogenetic tree of bacterial CsiD homologs. Homologs used in *in vitro* assays are highlighted in red. C) *in vitro* reactions of CsiD with different substrates using 2KG as a 2-oxoacid. Bars show peak area of 2-hydroxyacid, error bars show 95% CI, n=3. D) *In vitro* reactions of CsiD homologs with different 2-oxoacids. Bars represent spectral counts of L-2HG. Error bars show 95% CI, n=3.

Because non-heme Fe(II) oxidases can be promiscuous with respect to the 2-oxoacid cosubstrate (21, 22), we evaluated the 2-oxoacid specificity of CsiD. First, we evaluated the hydroxyl acceptor substrate specificity of CsiD family proteins by purifying two additional homologs from *E. coli* and a halophilic bacterium*, Halobacillus* sp. BAB-2008 (Figure 6b). We probed the activity of the homologs against a panel of 3 to 6 carbon fatty acids and diacids in the presence of 2KG, and found only glutarate served as a hydroxylation substrate (Figure 6c). These results are consistent with the work recently reported by Zhang *et al (9)* and further suggests the specificity of CsiD homologs is conserved across phyla. Although extremely specific for the hydroxylation substrate, all three CsiD homologs could utilize both 2OA and 2KG, but not oxaloacetate, as a cosubstrate for L-2HG formation (Figure 6d). The coproduct of the reaction using 2OA as a 2-oxoacid would be glutarate, rather than succinate. This result is particularly interesting as it provides a possible mechanism for the previously observed metabolic link between D-lysine and L-lysine catabolism. Growth defects observed in a ΔPP_2909 ΔPP_0158 double mutant grown on D-lysine also support this hypothesis (Figure S7a).

### Expression of lysine metabolic proteins is responsive to pathway metabolites

Multiple studies have demonstrated the expression of lysine catabolic genes is upregulated in the presence of pathway metabolites (9, 12, 23). To investigate the regulation of the newly-discovered lysine catabolic enzymes from this study, wild-type *P. putida* KT2440 was grown in minimal media on glucose or a single lysine metabolite (e.g. D-lysine, L-lysine, 5AVA, 2AA, or glutarate) as a sole carbon source until cultures reached an OD_600_ of 1.0. We then quantified the relative abundance of D- and L-lysine catabolic proteins via targeted proteomics (Figure 7). For each protein, all pairwise statistical comparisons of different carbon sources can be found in Supplemental Table 2. All five D-lysine pathway proteins measured (AmaA (PP_5257), AmaB (PP_5258), PP_4108, YdcJ (PP_5260), and YdiJ (PP_4493)) were upregulated when grown on L-lysine, D-lysine or 2AA compared to the glucose control. Neither 5AVA nor glutarate significantly induced expression of any measured D-lysine proteins. Of all the targeted proteins, the three identified in this study that directly degrade 2AA were most strongly induced by 2AA. Somewhat surprisingly, we also found the two enzymes involved in 2AA formation, AmaA and AmaB, were also more highly expressed in the presence of 2AA suggesting the possible involvement of a global regulator. An interesting finding from our initial screen showed sigma factor RpoX (PP_2088) to be required for fitness on D-lysine (Figure 1a). Deletion mutants of *rpoX* were severely attenuated in their ability to grown on D-lysine as a sole carbon source (Figure S7b). Further work will be necessary to examine complex regulatory network that controls D-lysine metabolism.

**Figure 7:**
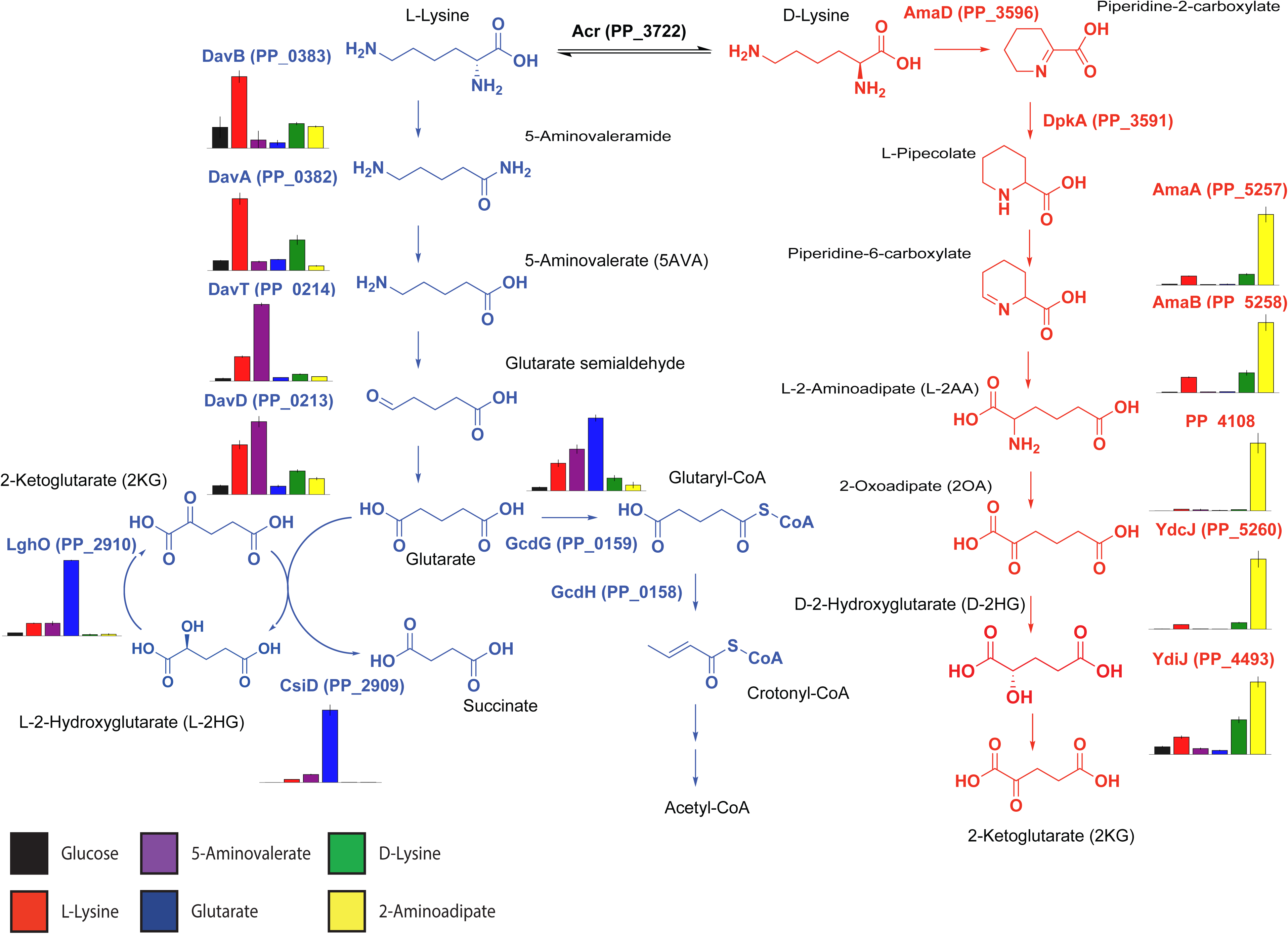
Expression of lysine degradation pathways in response to different lysine metabolites. Relative abundance of selected lysine degradation enzymes expressed in wild-type KT2440 in response to different carbon sources. Bars show spectral counts of proteins after 36 hours of growth on 10 mM glucose (black), 5AVA (purple), D-lysine (green), L-lysine (red), glutarate (blue), or 2AA (yellow). Error bars show 95% CI, n=3.

The initial two enzymes from L-lysine metabolism, DavA and DavB, were most highly expressed in the presence of L-lysine, but also significantly with D-lysine. As previously observed, DavT and DavD were most strongly upregulated on 5AVA, moderately upregulated on L-lysine, and to a lesser extent D-lysine. The induction of LhgO and CsiD was highest when grown on glutarate, although these proteins were also moderately upregulated by 5AVA and L-lysine. By comparison, PP_0159 (GcdG) expression in the presence of glutarate was stimulated to a lesser extent than LhgO and CsiD expression; in addition, GcdG was slightly upregulated on 5AVA and L-lysine.

## Discussion

Despite intensive study, a complete biochemical and genetic understanding of D-lysine catabolism in *P. putida* has remained elusive. A 2OA degradation pathway has been extensively characterized in mammals, because of its implications in human disease (24). In the mammalian pathway, L-lysine is metabolized to 2OA and eventually converted to acetyl-CoA via a glutaryl-CoA intermediate (24). However, this pathway has not been observed in bacteria. Previous work suggested 2OA is either converted via decarboxylation to glutarate or through several enzymatic steps to 2HG, yet none of these studies conclusively demonstrated a genetic and biochemical basis for these hypotheses (11, 19, 25). In this work we demonstrate plausible biochemical routes to account for both of these previously hypothesized pathways.

The first route, catalyzed by the DUF1338-containing metalloenzyme PP_5260, involves the direct conversion of 2OA to D-2HG. The formation of the D-2HG isomer by PP_5260 maintains stereochemical separation from the L-2HG formed by L-lysine degradation, thus requiring the dehydrogenase PP_4493 rather than the L-2HG specific oxidase *lghO*. This transformation seemingly involves two separate reactions: a decarboxylation and a hydroxylation. Hydroxymandelate synthase has been shown to catalyze a similar enzymatic reaction, via an intramolecular oxidative decarboxylation similar to 2KG dependent Fe(II) oxidases (26). PP_5260 is also a Fe(II) dependent decarboxylase, and the two share similar Kcat values for their given substrates (330 m^-1^ for PP_5260 and 270 m^-1^ for hydroxymandelate synthase) (27). Though PP_5260 and hydroxymandelate synthase share little sequence homology, this enzyme may give us insight into the molecular mechanism of DUF1338 enzymes. We have given PP_5260 the tentative title of 2-hydroxyglutarate synthase (*hglS*) until further mechanistic studies (currently underway in our group) are completed and a more accurate enzyme name can be assigned.

In bacteria, homologs of PP_5260 appear widely distributed with their genomic contexts suggesting functions both within and beyond lysine metabolism. Genomic contexts in other bacteria, particularly *Actinobacteria*, suggest these homologs may be involved in other amino acid catabolic pathways. Unfortunately, there is scant evidence for homologous function in model organisms. For example, although DUF1338 proteins are present in other *Ascomycota*, there is no homolog in *Saccharomyces cerevisiae*. Interestingly, the *E. coli* homolog of PP_5260 is located next to a potential glucans biosynthesis gene: Glucans biosynthesis protein D (28). Another DUF1338-containing protein from rice has been characterized and was implicated in starch granule formation (29). These results suggest DUF1338 proteins could play a role in sugar metabolism.

Recently Zhang *et al*. thoroughly characterized a CoA independent glutarate catabolism route ending at succinate involving the Fe(II) dependent oxygenase CsiD (9). Our RB-TnSeq screening convergently uncovered this pathway, and our biochemical and physiological results fully corroborate their findings. While both works show multiple CsiD homologs from divergent bacteria are highly specific towards glutarate as a hydroxyl acceptor, all three homologs we tested showed promiscuous activity toward 2-oxoacid cosubstrates. The ability of the *P. putida* CsiD to utilize 2OA as a cosubstrate is particularly interesting as it may directly connect L- and D-lysine metabolism. The promiscuity of CsiD may explain earlier studies which reported glutarate formation from D-lysine (11). Further studies involving labelled substrates may help elucidate the potential link between the two pathways. While CsiD plays a clear role in L-lysine metabolism in *P. putida*, its role in other organisms remains a mystery. In *E. coli*, RpoS controls the expression of CsiD, but *rpoS* mutants showed fitness benefits on all three lysine metabolites tested in our RB-TnSeq data (30). Recent work has shown that *E. coli* also uses CsiD to metabolize lysine, suggesting a possible conserved role for this pathway across bacteria (31).

Work presented here and previous reports have shown expression of both lysine catabolism pathways are highly responsive to their respective metabolites. While this metabolism appears highly coordinated, the genes themselves are dispersed across the genome, with both PP_4018, and PP_4493 found outside of operons, and relatively distant from other lysine catabolic genes. At least two global regulators appeared to be important to lysine metabolism based on our Rb-TnSeq data, *cbrA* (PP_4695) and *rpoX* (PP_2088). The two-component system CbrAB has been implicated in catabolite repression and C/N balance in *P. aeruginosa*, with mutants unable to grown on multiple amino acids (32). Further work in *P. putida* KT2440 showed the CbrAB system behaved similarly to that in *P. aeruginosa* (33). It would be unsurprising if this regulator controls the expression of various genes within lysine catabolism; more work into uncovering the regulon is warranted. RpoX on the other hand has been implicated in osmotic tolerance in *P. aeruginosa* (34, 35). This is interesting as lysine metabolism, and specifically pipecolate metabolism, has been associated with osmotic tolerance across multiple bacteria (36). As a *rpoX* deletion mutant was unable to grow on D-lysine, these results suggest D-lysine metabolism of *P. putida* may be involved in adaptation to saline or other osmotically stressful environments.

An interesting question remains as to why *P. putida* maintains separate metabolic pathways for D- and L-lysine, and why L-lysine metabolism seems dependent on the presence of an intact D-lysine metabolism. Previously work has proposed that the D-lysine pathway may provide a way of resolving a C/N imbalance that may occur when L-lysine is metabolized. However we believe this is unlikely as both lysine degradation pathways contain one deamination and one transamination reaction (11). Our fitness results indicate D-lysine metabolism is dispensable for growing on 5AVA. This would suggest only the initial two steps of L-lysine metabolism, the oxidation of lysine to 5-aminopentanamide by DavB and its subsequent deamination to 5AVA by DavA are dependent on D-lysine catabolism. We propose the adjacent AsnC family regulator PP_0384 likely responds to L-lysine as many proteins within this family respond to amino acids including lysine (37, 38) and expression of these two enzymes is most responsive to L-lysine. To our knowledge there has been no rigorous characterization of the regulation of the *davAB* operon, nor of the biochemical activities of these two enzymes *in vitro*. Future studies to uncover the mechanistic regulation at the transcriptional and post-translational levels at these two steps may uncover the necessity of D-lysine dependency of the L-lysine catabolic pathway. Overall our work highlights the utility of global fitness profiling to discover novel, complex, metabolic pathways in even well-characterized bacteria.

## Materials and Methods

### Media, chemicals, and culture conditions

Routine bacterial cultures were grown in in Luria-Bertani (LB) Miller medium (BD Biosciences, USA). *E. coli* was grown at 37 °C, while *P. putida* was grown at 30 °C unless otherwise noted. When indicated, *P. putida* was grown on modified MOPS minimal medium (39). Cultures were supplemented with kanamycin (50 mg/L, Sigma Aldrich, USA), gentamicin (30 mg/L, Fisher Scientific, USA), or carbenicillin (100mg/L, Sigma Aldrich, USA), when indicated. D-2AA was purchased from Takara Bioscience (USA), all other compounds were purchased through Sigma Aldrich.

### Strains and plasmids

All bacterial strains and plasmids used in this work are listed in Supplemental Table 1. All strains and plasmids created in this work are available through the public instance of the JBEI registry. (https://public-registry.jbei.org/folders/391). All plasmids were designed using Device Editor and Vector Editor software, while all primers used for the construction of plasmids were designed using j5 software (40–42). Synthetic DNA coding for the *Halobacillus* sp. BAB-2008 *csiD* homolog was purchased from Integrated DNA Technologies (IDT, Coralville, IA). Plasmids were assembled via Gibson Assembly using standard protocols (43), or Golden Gate Assembly using standard protocols (44). Plasmids were routinely isolated using the Qiaprep Spin Miniprep kit (Qiagen, USA), and all primers were purchased from Integrated DNA Technologies (IDT, Coralville, IA).

### Random barcode TnSeq experiments

*P. putida* RB-TnSeq library JBEI-1 was created by diluting a 1 mL aliquot of the previously described *P. putida* RB-TnSeq library (16) in 500 mL of LB media supplemented with kanamycin which was then grown to an OD_600_ of 0.5 and frozen as 1 mL aliquots after adding glycerol to a final concentration of 20% v/v. Libraries were stored at −80 °C until used. A 1 mL aliquot of *P. putida* RB-TnSeq library JBEI-1 was thawed on ice and diluted in 25 mL of LB supplemented with kanamycin. The culture was grown until it reached an OD_600_ of 0.5, at which point 3 1-mL aliquots were taken, pelleted, decanted, and then stored at −80 °C to use as a time zero control. The library was then washed once in MOPS minimal medium without any carbon source, and then diluted 1:50 into 10 mL MOPS minimal medium supplemented with either 10 mM glucose, 5AVA, D-lysine, or L-lysine. Cells were grown in 50 mL culture tubes for 48 hours at 30 °C shaking at 200 rpm. After growth 2 ml aliquots from the culture tubes were pelleted, decanted and frozen at −80 °C for barcode sequencing. We performed DNA barcode sequencing (BarSeq) as previously described (14, 16). The fitness of a strain is the normalized log2 ratio of barcode reads in the experimental sample to barcode reads in the time zero sample. The fitness of a gene is the weighted average of the strain fitness for insertions in the central 10– 90% of the gene. The gene fitness values are normalized so the typical gene has a fitness of zero. The primary statistic *t*-value is of the form of fitness divided by the estimated variance across different mutants of the same gene. Statistic *t*-values of >|4| were considered significant. All experiments described herein pass the quality metrics described previously unless noted otherwise. All fitness data in this work is publically available at http://fit.genomics.lbl.gov.

### Construction of deletion mutants

Deletion mutants in *P. putida* were constructed by homologous recombination and *sacB* counterselection using the allelic exchange vector pMQ30 (45). Briefly, homology fragments ranging from 1 kbp to 2 kbp up- and downstream of the target gene, including the start and stop codons respectively, were cloned into pMQ30. An exception to these design parameters was plasmid pMQ-PP_5260 which maintained an additional 21 nt at the 5’ end of the gene in addition to the stop codon. Plasmids were then transformed via electroporation into *E. coli* S17 and then mated into *P. putida* via conjugation. Transconjugants were selected for on LB agar plates supplemented with gentamicin 30 mg/mL, and chloramphenicol 30 mg/mL. Transconjugants were then grown overnight on LB media also supplemented with 30 mg/mL gentamicin, and 30 mg/mL chloramphenicol, and then plated on LB agar with no NaCl supplemented with 10% w/v sucrose. Putative deletions were restreaked on LB agar with no NaCl supplemented with 10% w/v sucrose, and then were screened via PCR with primers flanking the target gene to confirm gene deletion.

### Plate based growth assays

Growth studies of bacterial strains were conducted a microplate reader kinetic assays. Overnight cultures were inoculated into 10 mL of LB medium from single colonies, and grown at 30 °C. These cultures were then washed twice with MOPS minimal media without any added carbon and diluted 1:100 into 500 uL of MOPS medium with 10 mM of a carbon source in 48-well plates (Falcon, 353072). Plates were sealed with a gas-permeable microplate adhesive film (VWR, USA), and then optical density was monitored for 48 hours in an Biotek Synergy 4 plate reader (BioTek, USA) at 30 °C with fast continuous shaking. Optical density was measured at 600 nm and all OD_600_ measurements are reported without pathlength corrections.

### Expression and purification of proteins

A 5 mL overnight culture of *E. coli* BL21 (DE3) containing the expression plasmid was used to inoculate a 500 mL culture of LB. Cells were grown at 37 °C to an OD_600_ of 0.6 then induced with isopropyl β-D-1-thiogalactopyranoside to a final concentration of 1 mM. The temperature was lowered to 30 °C and cells were allowed to express for 18 hours before being harvested via centrifugation. Cell pellets were stored at −80 °C until purification. For purification, cell pellets were resuspended in lysis buffer (50 mM sodium phosphate, 300 mM sodium chloride, 10 mM imidazole, 8% glycerol, pH 7.5) and sonicated to lyse cells. Insolubles were pelleted via centrifugation (30 minutes at 40,000xg). The supernatant was applied to a fritted column containing Ni-NTA resin (Qiagen, USA) which had been pre-equilibrated with several column volumes of lysis buffer. The resin was washed with lysis buffer containing 50 mM imidazole, then the protein was eluted using a step-wise gradient of lysis buffer containing increasing imidazole concentrations (100 mM, 200 mM, and 400 mM). Fractions were collected and analyzed via SDS-PAGE. Purified protein was dialyzed overnight at 4 °C against 50 mM HEPES pH 7.5, 5% glycerol.

### CsiD *in vitro* assays

The activity of purified CsiD homologs was analyzed in 100 µL reaction mixtures containing 50 mM HEPES (pH 7), 5 mM glutarate, 5 mM 2KG, 25 µM FeSO_4_, 0.1 mM ascorbate, and 0.5 mM dithiothreitol. For negative control reactions, each respective reaction component was omitted. To initiate reactions, CsiD was added to a final concentration of 7 µM. For the no enzyme control, CsiD was denatured at 98 °C for 10 minutes prior to addition to the reaction mix. Reactions were allowed to proceed at 22 °C for 3 hours. Products were analyzed via LC-TOF method 3 after quenching via the addition of acetonitrile and methanol for a final ACN:H_2_O:MeOH ratio of 6:3:1 To analyze products from substrate range as well 2-oxoacid specificity experiments, reactions were measured via LC-TOF method 1.

### Transamination assays

To determine product formation via PP_4108, assays were conducted in 50 mM HEPES (pH 7.5), with 5 mM 2KG, 0.1 mM PLP, and 5 mM of substrate, and 10 µM of purified enzyme or boiled enzyme control in 100µL volumes. Reactions were incubated at 30 °C for 16 hours and quenched via the addition of ACN and MeOH for a final ACN:H_2_O:MeOH ratio of 6:3:1 for LC-TOF method 3. To determine substrate specificity reactions were set up at 75 µL scale and carried out at 30°C for up to 16 hours before freezing. For analysis, reactions were diluted 15-fold in water and assessed by a colorimetric assay for glutamate (Sigma MAK004) in 96-well format via a SpectraMax M4 plate reader (Molecular Devices, USA).

### PP_5260 *in vitro* assays

The activity of PP_5260 was initially assessed in 50 mM HEPES, with 5 mM 2OA as substrate and 10 µM purified enzyme or boiled enzyme control. Reactions were incubated for 16 hours at 30 °C. To test the necessity of metal cofactors EDTA was added to a final concentration of 50 µM. Reactions and quenched via the addition of ACN and methanol MeOH for a final ACN:H_2_O:MeOH ratio of 6:3:1 for LC-TOF analysis method 3, or with an equal volume of ice-cold methanol for HPLC analysis and LC-TOF method 2.

To determine the metal cofactor, after purification over Ni-NTA resin the protein was concentrated and dialyzed overnight against 50mM HEPES, 100mM NaCl, pH 7.5. To generate apo-enzyme, the protein was then dialyzed four times at a protein:dialysis buffer ratio of 1:300 against the same buffer containing 5mM EDTA in order to remove any bound metal. The enzyme was dialyzed once more against buffer without EDTA overnight in order to remove any remaining chelating reagent. The apo-enzyme was then assayed in the presence of 50 µM of a variety of potential metal cofactors in 50 mM HEPES with 10 mM 2OA as substrate and 10 µM purified enzyme. Reactions were incubated for 30 minutes at 30 °C and activity was assessed via HPLC.

Determination of enzyme stoichiometry was assessed in 50 mM HEPES, 50 µM FeCl_2_, with 1 mM 2OA as substrate and 0.1 µM purified enzyme or boiled enzyme control. Reactions were incubated for 5 minutes at 30 °C and then quenched with an equal volume of ice-cold methanol then quantified with LC-TOF method 2.

Enzyme coupled decarboxylation assays were carried out as previously described (20). Reaction mixtures contained 100 mM Tris-HCl (pH 7), 10 mM MgCl_2_, 0.4 mM NADH, 4 mM, 50 µM FeCl_2_, phosphoenol pyruvate (PEP), 100U/mL pig heart malate dehydrogenase(Roche), 2U/mL microbial PEP carboxylase (Sigma), and 10 mM 2OA. Reactions were initiated by the addition of purified PP_5260 or boiled enzyme controls, and absorbance at 340 nm was measured via a SpectraMax M4 plate reader (Molecular Devices, USA). Michaelis-Menten behavior was formulated as previously described (46). Least-squares minimization was used to derive K_m_ and K_cat_. Determination of D-2HG concentration was assayed with a D-2-Hydroxyglutarate (D-2HG) Assay Kit (Sigma MAK320).

**Table 1.**

| **Strain** | **JBEI Part ID** | **Reference** | **Genotype** |
| --- | --- | --- | --- |
| *E. coli* DH10B |  | ThermoFisher |  |
| *E. coli* S17 |  | ATCC 47055 |  |
| *E. coli* BL21(DE3) |  | Novagen |  |
| *P. putida* KT2440 |  | ATCC 47054 |  |
| *P. putida* ΔPP_0158 | JPUB_010967 | This work |  |
| *P. putida* ΔPP_2088 | JPUB_013224 | This work |  |
| *P. putida* ΔPP_2909 | JPUB_010968 | This work |  |
| *P. putida* ΔPP_2910 | JPUB_010969 | This work |  |
| *P. putida* ΔPP_0158<br>ΔPP_2909 | JPUB_010970 | This work |  |
| *P. putida* ΔPP_4108 | JPUB_010971 | This work |  |
| *P. putida* ΔPP_4493 | JPUB_010972 | This work |  |
| *P. putida* ΔPP_5260 | JPUB_010973 | This work |  |
| **Plasmids** |  |  |  |
| pMQ30 |  | 45 | Gm, SacB |
| pET28 |  | Novagen | Kan |
| pET21b |  | Novagen | Amp |
| pMQ30-PP_0158 | JPUB_010989 | This work | Gm, SacB |
| pMQ30-PP_2088 | JPUB_013222 | This work | Gm, SacB |
| pMQ30-PP_2909 | JPUB_010991 | This work | Gm, SacB |
| pMQ30-PP_2910 | JPUB_010995 | This work | Gm, SacB |
| pMQ30-PP_4108 | JPUB_010981 | This work | Gm, SacB |
| pMQ30-PP_4493 | JPUB_010979 | This work | Gm, SacB |
| pMQ30-PP_5260 | JPUB_010977 | This work | Gm, SacB |
| pET28-CsiD_Halo | JPUB_010987 | This work | Kan |
| pET28-CsiD_Ecoli | JPUB_010993 | This work | Kan |
| pET28-CsiD_Pput | JPUB_010975 | This work | Kan |
| pET21b-PP_4108 | JPUB_010983 | This work | Amp |
| pET21b-PP_5260 | JPUB_010985 | This work | Amp |

## Acknowledgements

This manuscript is dedicated to the memory of Cornell Professor Dr. Eugene Madsen. The authors would like to thank Morgan Price, Dr. John Hangasky, Dr. Jamie Meadows, Dr. Robert Haushalter, Dr. Bo Pang, Dr. Nick Weathersby, Mary Thompson, and Catharine Adams for their helpful discussions in preparing this manuscript. We would also like to thank the UC Berkeley SMART program for providing support for R.K to conduct summer research. This work was part of the DOE Joint BioEnergy Institute (https://www.jbei.org) supported by the U.S. Department of Energy, Office of Science, Office of Biological and Environmental Research, and protein purification and homology modelling components were part of the Agile BioFoundry (http://agilebiofoundry.org) supported by the U.S. Department of Energy, Energy Efficiency and Renewable Energy, Bioenergy Technologies Office, through contract DE-AC02-05CH11231 between Lawrence Berkeley National Laboratory and the U.S. Department of Energy. The views and opinions of the authors expressed herein do not necessarily state or reflect those of the United States Government or any agency thereof. Neither the United States Government nor any agency thereof, nor any of their employees, makes any warranty, expressed or implied, or assumes any legal liability or responsibility for the accuracy, completeness, or usefulness of any information, apparatus, product, or process disclosed, or represents that its use would not infringe privately owned rights.

## Contributions

Conceptualization, M.G.T., and J.M.B.; Methodology, M.G.T., J.M.B, J.F.B., P.C.M., S.C.C., N.C.H, C.B.E, E.E.K.B, C.J.P., and A.M.D.; Investigation, M.G.T., J.M.B, W.N.S., R.A.K, J.F.B., V.T.B, P.C.M., J.W.G, C.J.P, N.C.H., F.F.T., J.H.P W.S., E.E.K.B.; Writing – Original Draft, M.G.T.; Writing – Review and Editing, All authors.; Resources and supervision, P.D.A., A.P.A., A.M.D., and J.D.K.

## Supplementary Materials and Methods

### HPLC analysis

HPLC analysis was performed on an Agilent Technologies 1200 series liquid chromatography instrument coupled to a refractive index detector (35°C, Agilent Technologies, Santa Clara, CA). Samples were injected onto an Aminex HPX-87H Ion Exclusion Column (300 x 7.8 mm, 60°C, Bio-Rad, Hercules, CA) and eluted isocratically with 4 mM H_2_SO_4_ at 600 uL/min for 20 minutes. Compounds were quantified via comparison to a calibration curve prepared with authentic standards and normalized to injection volume.

### Proteomics analysis

*P. putida* KT2440 wild type was grown on MOPS minimal media with 10 mM of either glucose, L-lysine, D-lysine, 5AVA, 2AA, or glutarate. Cells were harvested when cultures reached an OD_600_ of 1.0 with a 1 cm pathlength. Cell lysis and protein precipitation were achieved by using a chloroform-methanol extraction as previously described (1). Thawed pellets were loosened from 14 mL falcon tubes and transferred to PCR 8-well tube strip, followed by the addition of 80 µL of methanol, 20 µL of chloroform, and 60 µL of water, with vortexing. The samples were centrifuged at ∼20,000 x g for 1 minute for phase separation. The methanol and water (top) layer was removed, then 100 µL of methanol was added and the sample was vortexed briefly. The samples were centrifuged at ∼20,000 x g for 1 minute to isolate the protein pellet. The protein pellet was air-dried for 10 minutes and resuspended in 100 mM ammonium bicarbonate with 20% methanol. The protein concentration was measured using the DC Protein Assay Kit (Bio-Rad, Hercules, CA) with bovine serum albumin for the standard curve. A total of 100 µg of protein from each sample was digested with trypsin for targeted proteomic analysis. The protein was reduced by adding tris 2-(carboxyethyl) phosphine (TCEP) at a final concentration of 5 mM, alkylated by adding iodoacetamide at a final concentration of 10 mM, and digested overnight at 37 °C with trypsin at a ratio of 1:50 (w/w) trypsin:total protein. As previously described (2), peptides were analyzed using an Agilent 1290 liquid chromatography system coupled to an Agilent 6460QQQ mass spectrometer (Agilent Technologies, Santa Clara, CA). Peptide samples (10 µg) were separated on an Ascentis Express Peptide ES-C18 column (2.7 µm particle size, 160 Å pore size, 50 mm length x 2.1 mm i.d., 60 °C; Sigma-Aldrich, St. Louis, MO) by using a chromatographic gradient (400 µL/min flow rate) with an initial condition of 95% buffer A (99.9% water, 0.1% formic acid) and 5% buffer B (99.9% acetonitrile, 0.1% formic acid) then increasing linearly to 65% buffer A/35% buffer B over 5.5 minutes. Buffer B was then increased to 80% over 0.3 minutes and held at 80% for two minutes followed by ramping back down to 5% buffer B over 0.5 minutes where it was held for 1.5 minutes to re-equilibrate the column for the next sample. The peptides were ionized by an Agilent Jet Stream ESI source operating in positive-ion mode with the following source parameters: gas Temperature: 250 °C, gas Flow: 13 L/min, nebulizer pressure: 35 psi, sheath gas temperature: 250 °C, sheath gas flow: 11 L/min, nozzle voltage: 0 V, chamber voltage: 3,500 V. The data were acquired using Agilent MassHunter, version B.08.02, processed using Skyline (3) version 4.1, and peak quantification was refined with mProphet (4) in Skyline. Data are available at Panorama Public via this link: https://panoramaweb.org/massive_fitness_profiling_Pseudomonas_putida.url. All pairwise combinations of spectral counts from carbon sources for each protein were compared via Student’s t-test followed by a Bonferroni correction.

### Detection of metabolites

Sampling of intracellular metabolites was conducted as described previously (5). Multiple methods were used to detect compounds in this work. Method (1) HILIC-HRMS analysis was performed using an Agilent Technologies 6510 Accurate-Mass Q-TOF LC-MS instrument using positive mode and an Atlantis HILIC Silica 5 µm column (150 x 4.6 mm) with a linear of 95 to 50% acetonitrile (v/v) over 8 minutes in water with 40 mM ammonium formate, pH 4.5, at a flow rate of 1 mL minute^−1^. Method (2) HILIC-HRMS analysis was performed using an Agilent Technologies 6510 Accurate-Mass Q-TOF LC-MS instrument using negative mode and an Atlantis HILIC Silica 5 µm column (150 x 4.6 mm) with an isocratic mobile phase (80% acetonitrile (v/v) with 40 mM ammonium formate, pH 4.5) for 20 minute at a flow rate of 1 mL minute^−1^. Method (3) is described in George et al (5). Briefly, samples were separated via a SeQuantZIC-pHILIC guard column (20-mm length, 2.1-mm internal diameter, and 5-μm particle size; from EMD Millipore, Billerica, MA, USA), then with a short SeQuantZIC-pHILIC column (50-mm length, 2.1-mm internal diameter, and 5-μm particle size) followed by a long SeQuantZIC-pHILIC column (150-mm length, 2.1-mm internal diameter, and 5-μm particle size) using an Agilent Technologies 1200 Series Rapid Resolution HPLC system (Agilent Technologies, Santa Clara, CA, USA). The mobile phase was composed of 10 mM ammonium carbonate and 118.4 mM ammonium hydroxide in acetonitrile/water (60.2:39.8, v/v). Metabolites were eluted isocratically via a flow rate of 0.18 mL/min from 0 to 5.4 minutes, which was increased to 0.27 mL/min from 5.4 to 5.7 minutes, and held at this flow rate for an additional 5.4 minutes. The HPLC system was coupled to an Agilent Technologies 6210 TOF-MS system in negative mode. Determination of D-2HG concentration was assayed with a D-2-Hydroxyglutarate (D2HG) Assay Kit (Sigma MAK320).

### Phylogenomic analyses

Amino acid sequences of CsiD homologs were downloaded from the pFAM database and aligned with MAFFT-linsi (13). Phylogenetic trees of CsiD alignments were constructed with FastTree 2, and trees were visualized on iTOL (14, 15).

Representative DUF1338 sequences were obtained from pFAM (https://pfam.xfam.org/family/PF07063#tabview=tab3). All genomes analyzed were downloaded from the NCBI FTP site and annotated using RAST (16). Amino acid sequences of DUF1338 proteins from these genomes were retrieved using BlastP with a bit score cutoff of 150 and an E-value of 0.000001. All sequences alignments were performed using Muscle v3.8 (17) and the alignments were manually curated using Jalview V2 (18).

For the phylogenetic reconstructions, the best amino acid substitution model was selected using ModelFinder implemented on IQ-tree (19) the phylogenies were obtained using IQ-tree v 1.6.7 (20), with 10,000 bootstrap replicates. The final trees were visualized and annotated using FigTree v1.4.3 (http://tree.bio.ed.ac.uk/software/figtree/). Genome neighborhoods of DUF1338 were obtained using CORASON-BGC (21) and manually colored and annotated.

### Statistical analyses and data presentation

All numerical data were analyzed using custom Python scripts. All graphs were visualized using either Seaborn or Matplotlib. Calculation of 95% confidence intervals, standard deviations, and T-test statistics were conducted via the Scipy library. Bonferroni corrections were calculated using the MNE python library (22).

**Figure S1:**
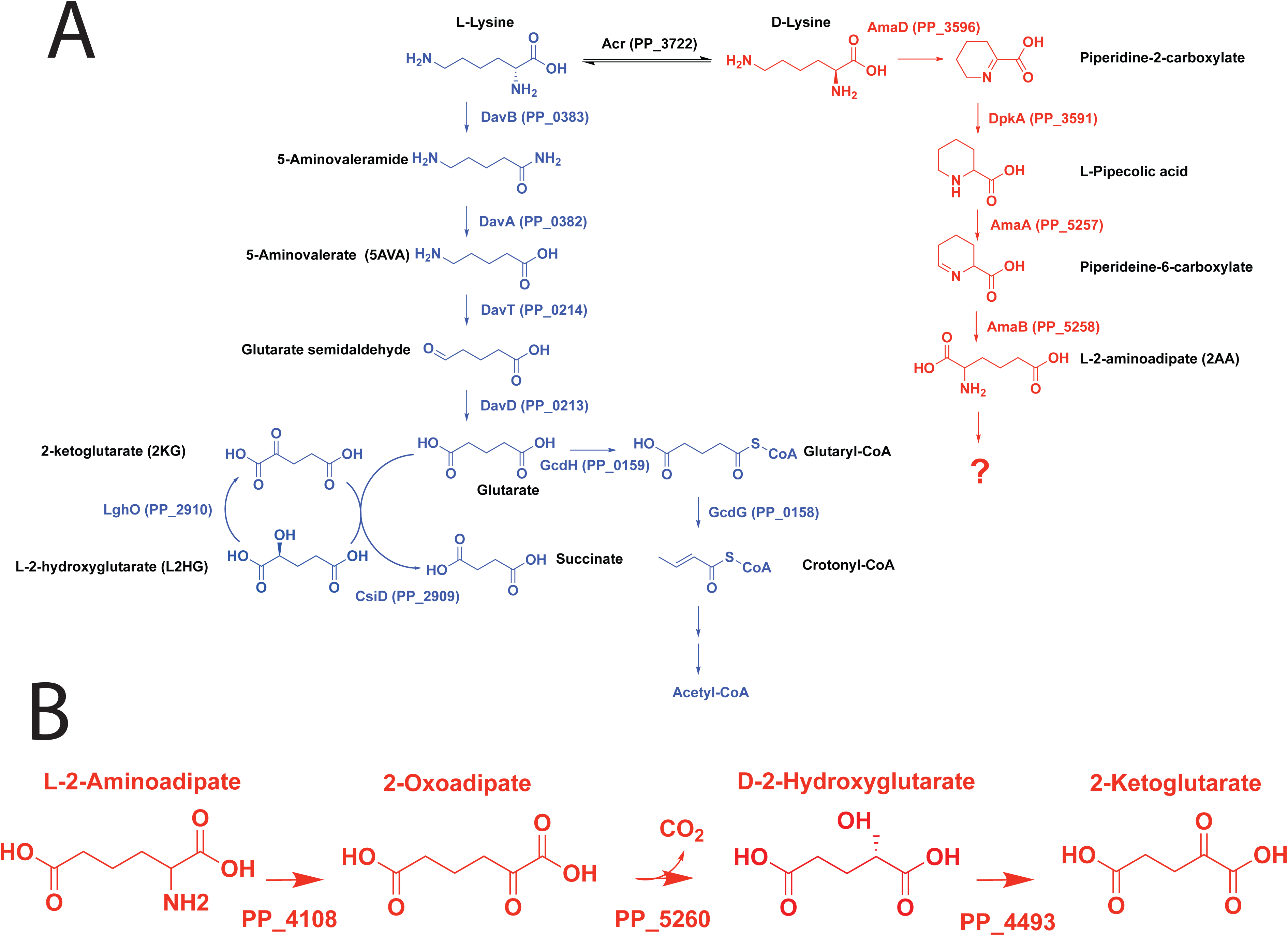
Metabolic pathways of lysine catabolism in *P. putida* KT2440. A) L-lysine metabolic pathway is shown in blue, while the known steps of D-lysine metabolism are shown in red. B) Proposed route of 2AA metabolism in *P. putida*.

**Figure S2:**
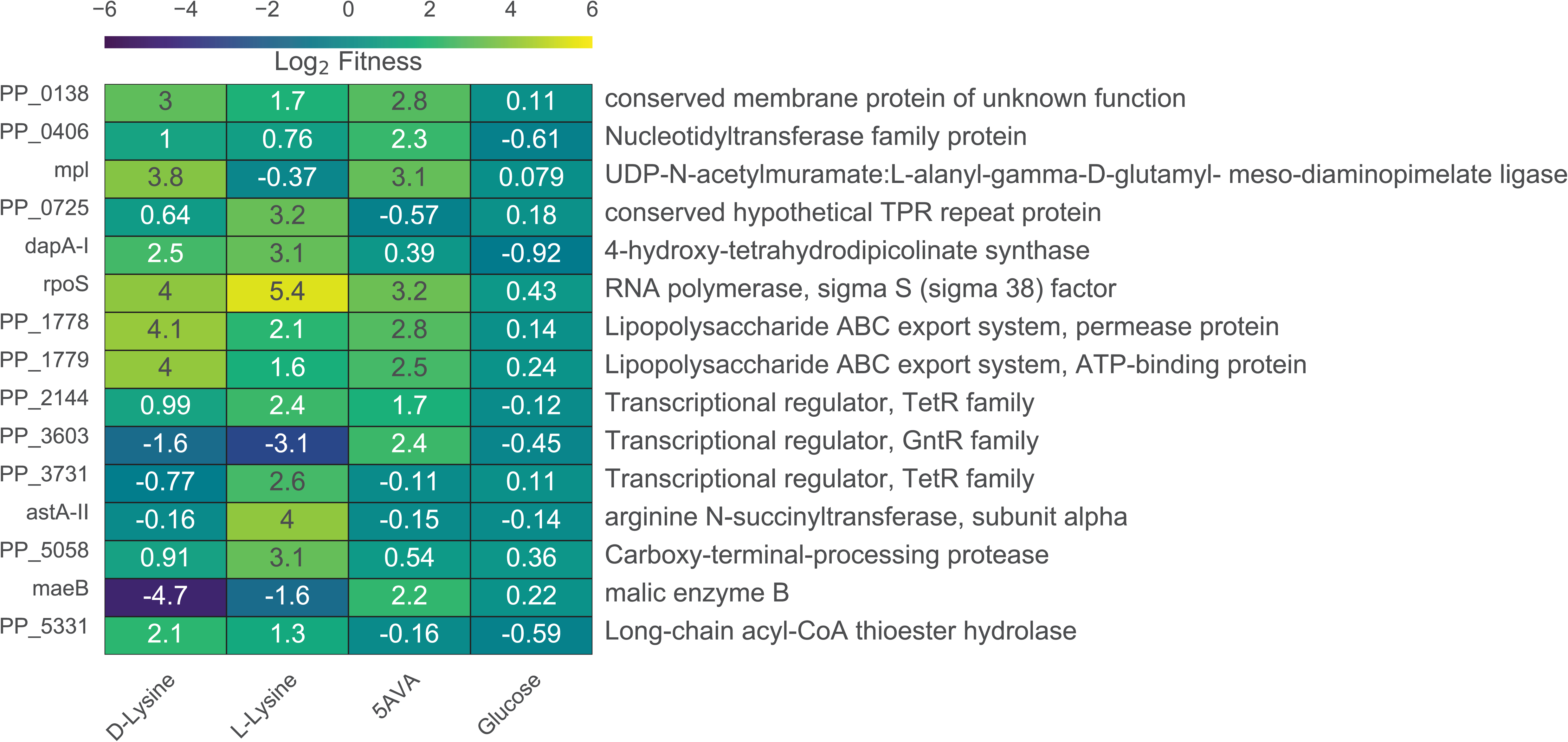
Results of RB-TnSeq screen. A) Genes that showed great than 2 log2 fitness on either D-lysine, L-lysine, or 5AVA but showed no less than 0.5 log2 fitness defect when grown on glucose.

**Figure S3:**
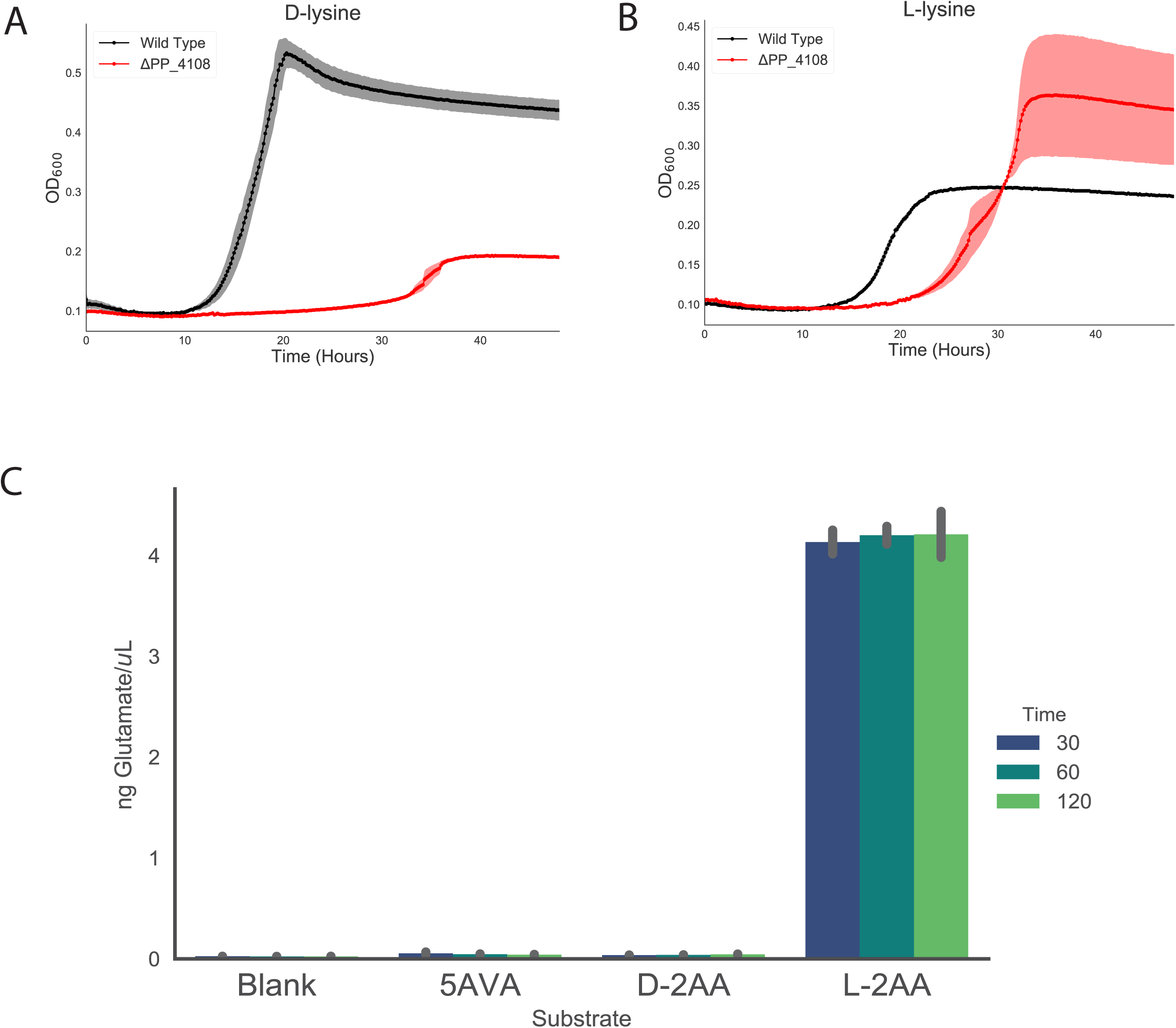
Growth of PP_4108 mutants on lysine. A) Growth of wild-type KT2440 and PP_4108 mutant on D-lysine as a sole carbon source. Shaded area represents 95% CI, n=3. B) Growth of wild-type KT2440 and PP_4108 mutant on L-lysine as a sole carbon source. Shaded area represents 95% CI, n=3. C) Colorimetric glutamate formation time course of PP_4108. Error bars are standard deviation of n=2.

**Figure S4:**
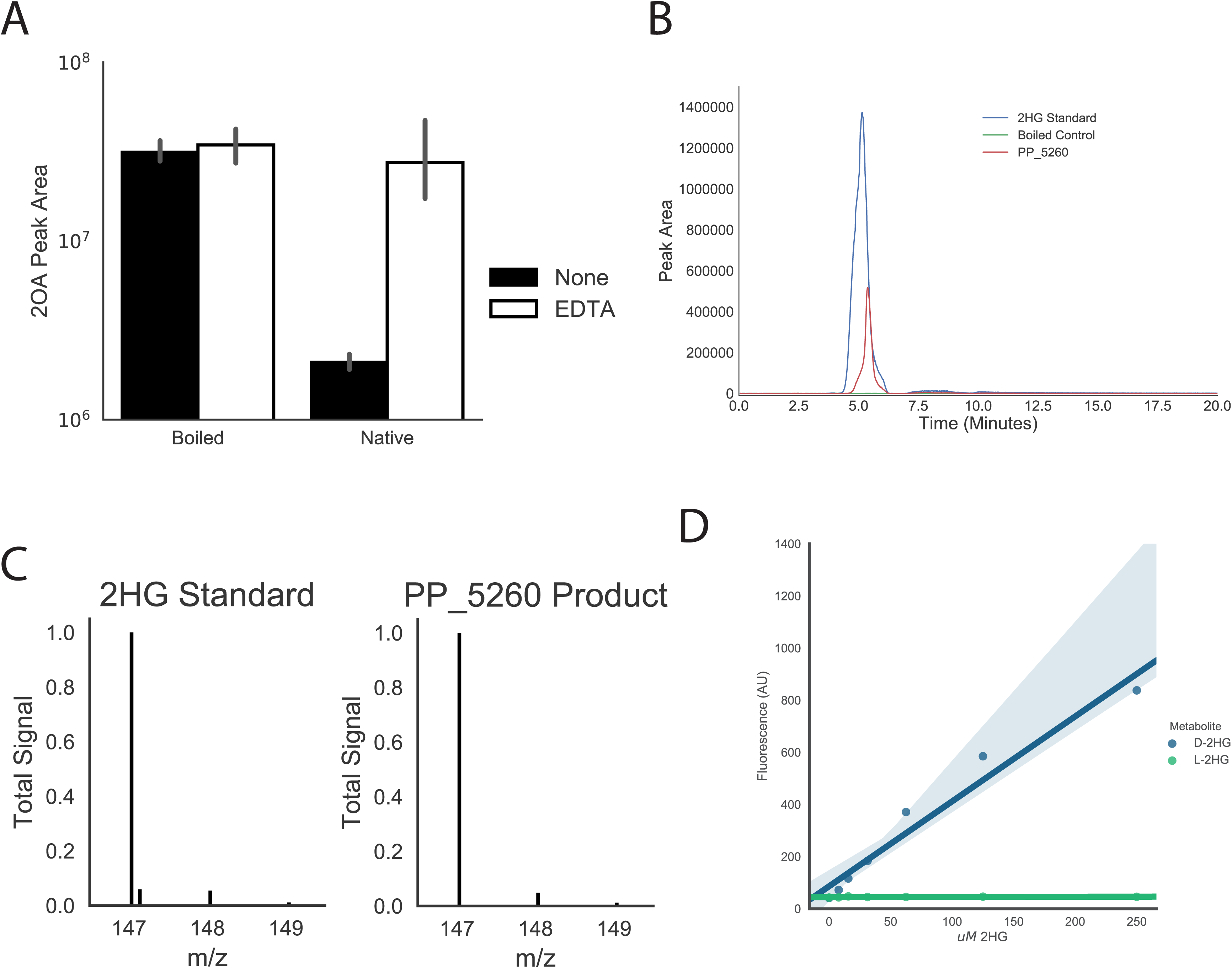
Characterization of PP_5260. A) In vitro reactions of boiled or native PP_5260 incubated with 2OA with 50 µM EDTA (white), or without EDTA (black). Bars represent 1og_10_ transformed spectral counts, errors bars represent 95% CI, n=3 B) E LC-TOF analysis of a 2HG standard, and products of PP_5260 incubated with 2OA, and boiled control. C) Mass spectra of 2HG standard and product of PP_5260 in vitro reaction. D) Standard curves of D-2HG and L-2HG using a D-2HG specific enzymatic detection assay. Shaded areas represent 95% CI, n=3.

**Figure S5:**
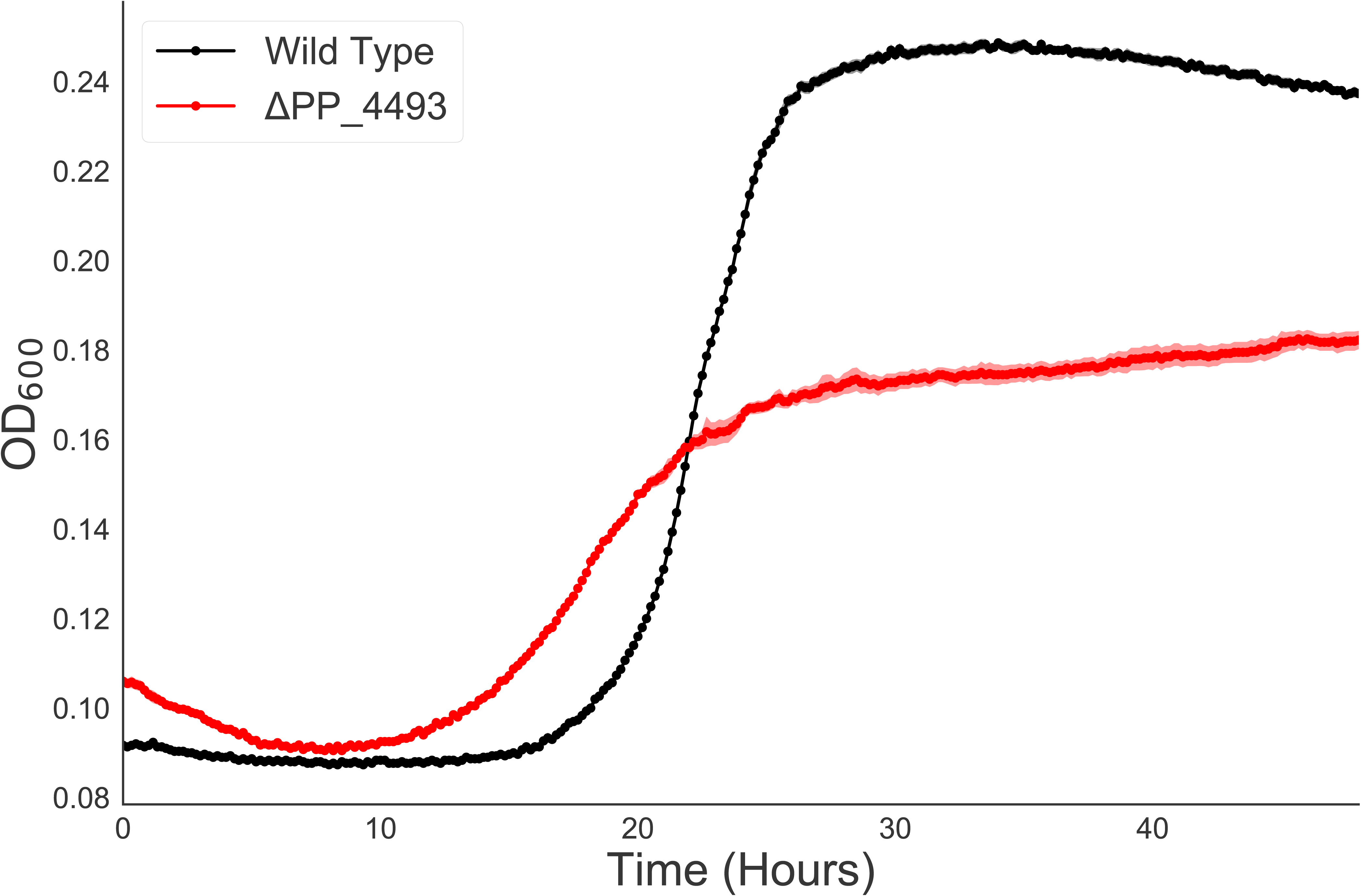
Growth of PP_4493 mutant on L-lysine. Growth of wild-type KT2440 and PP_4493 mutant on L-lysine as a sole carbon source. Shaded area represents 95% CI, n=3

**Figure S6:**
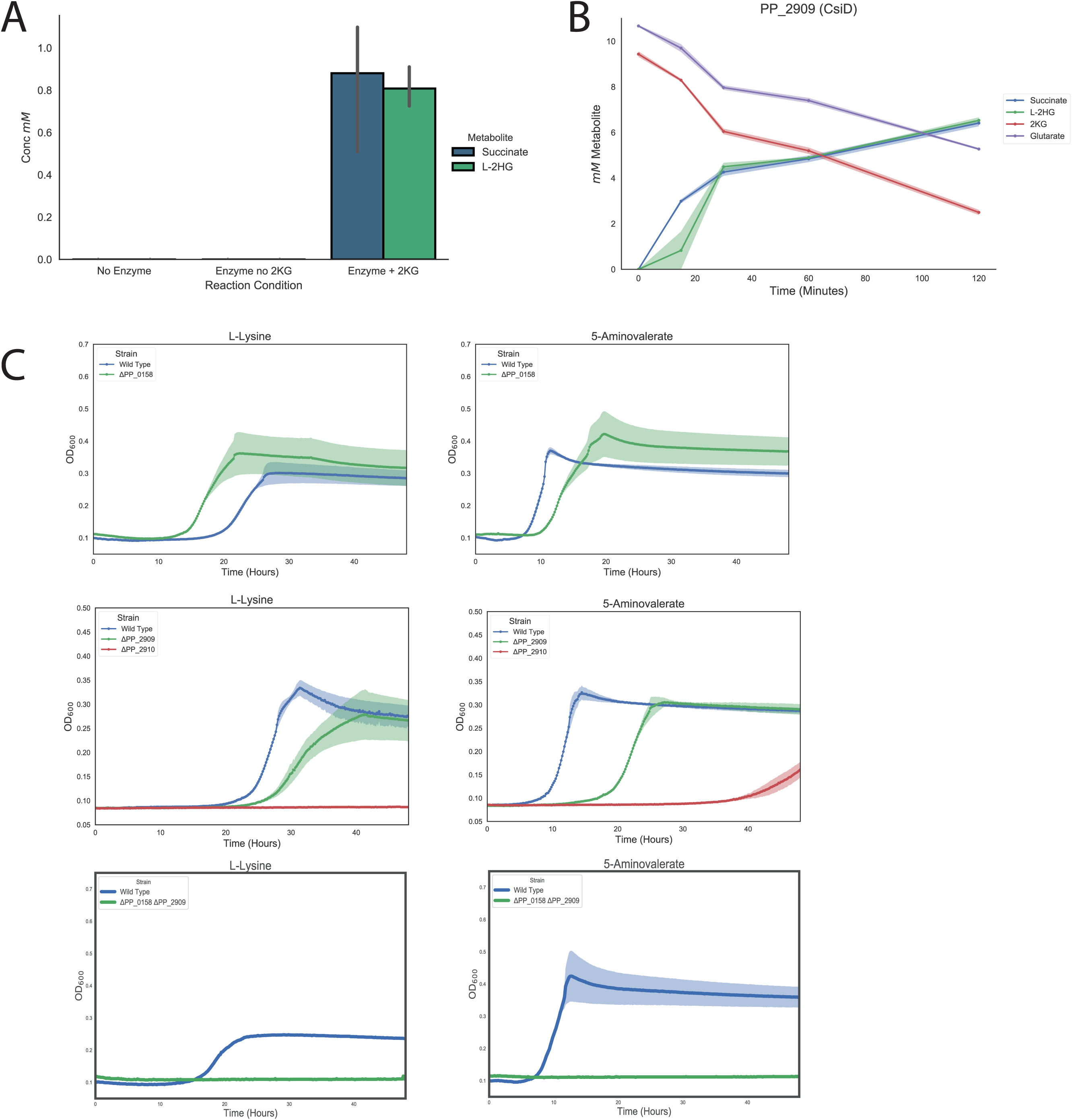
Characterization of PP_2909 (CsiD) in *P. putida*. A) In vitro reactions of PP_2909. Bars show mM of either succinate (blue), or 2HG (green) formed by boiled enzyme control, no 2KG control, or native enzyme with 2KG added. Errors bars show 95% CI, n=3. B) Time course in vitro reaction of PP_2909. Plot shows 2HG, 2KG, succinate, and glutarate overtime. Shaded region shows 95% CI, n=3. C) Growth curves of wild-type KT2440, PP_0158, PP_2909, PP_2910, or PP_2909/PP_0158 double mutants grown on either L-lysine (left column), or 5AVA (right column). Shaded region shows 95% CI, n=3.

**Figure S7:**
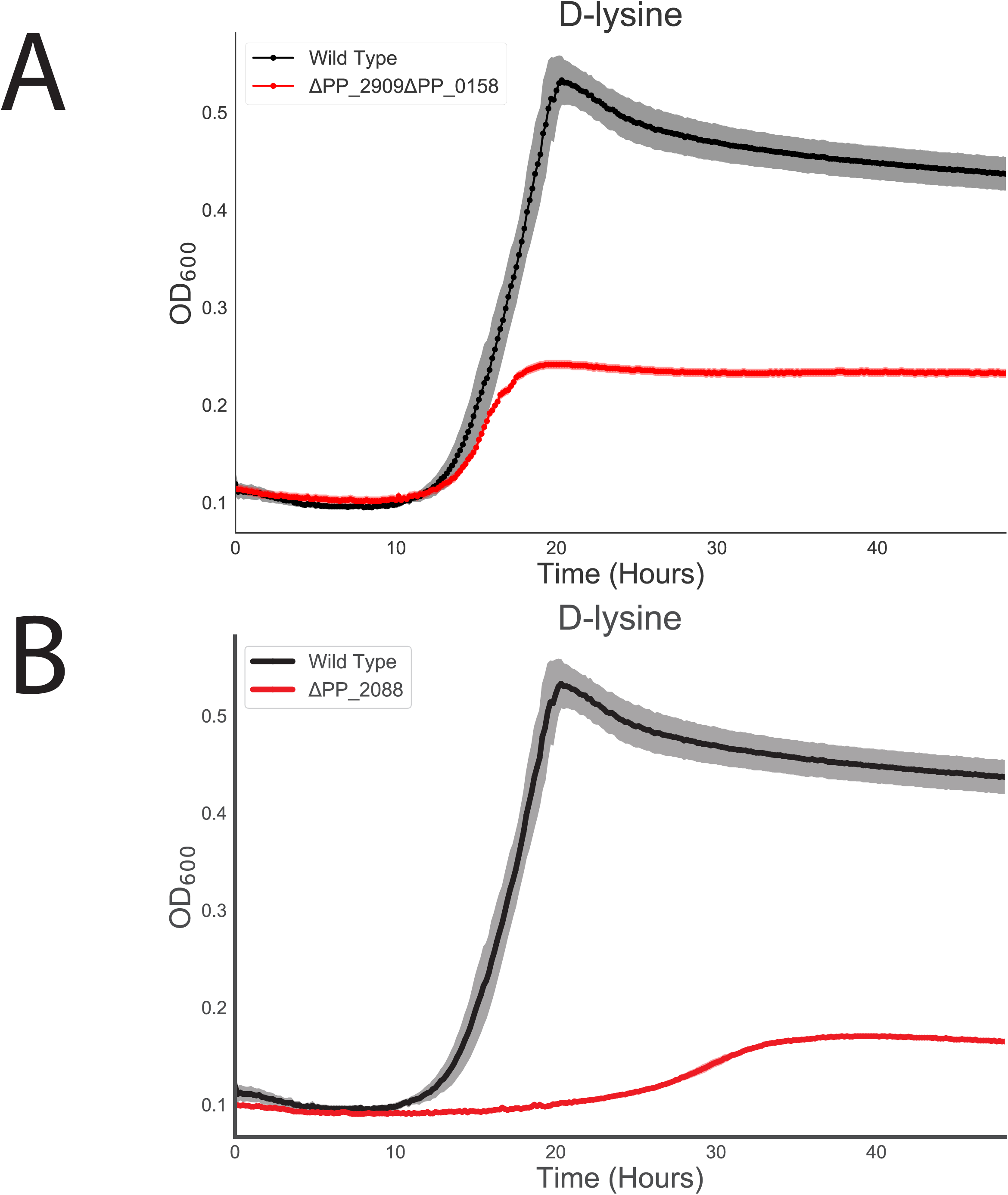
Growth of PP_2909/PP_0158 and PP_2088 mutants on D-lysine. A) Growth of wild-type KT2440 and PP_2909/PP_0158 mutant on D-lysine as a sole carbon source. Shaded area represents 95% CI, n=3. A) Growth of wild-type KT2440 and PP_2088 mutant on D-lysine as a sole carbon source. Shaded area represents 95% CI, n=3.

